# Expectation generation and its effect on subsequent pain and visual perception

**DOI:** 10.1101/2024.10.10.617570

**Authors:** Rotem Botvinik-Nezer, Stephan Geuter, Martin A. Lindquist, Tor D. Wager

## Abstract

Bayesian accounts of perception, such as predictive processing, suggest that perceptions integrate expectations and sensory experience, and thus assimilate to expected values. Furthermore, more precise expectations should have stronger influences on perception. We tested these hypotheses in a paradigm that manipulates both the mean value and the precision of cues within-person. Forty-five participants observed cues–presented as ratings from 10 previous participants–with varying cue means, variances (precision), and skewness across trials. Participants reported expectations regarding the painfulness of thermal stimuli or the visual contrast of flickering checkerboards. Subsequently, similar cues were each followed by a visual or noxious thermal stimulus. While perceptions assimilated to expected values in both modalities, cues’ precision mainly affected visual ratings. Furthermore, behavioral and computational models revealed that expectations were biased towards extreme values in both modalities, and towards low-pain cues specifically. fMRI analysis revealed that the cues affected systems related to higher-level affective and cognitive processes–including assimilation to the cue mean in a neuromarker of endogenous contributions to pain and in the nucleus accumbens, and activity consistent with aversive prediction-error-like encoding in the periaqueductal gray during pain perception–but not systems related to early perceptual processing. Our findings suggest that predictive processing theories should be combined with mechanisms such as selective attention to better fit empirical findings, and that expectation generation and its perceptual effects are mostly modality-specific and operate on higher-level processes rather than early perception.

## Introduction

Current theories of perception, such as Bayesian theories ^1,2^ and predictive processing ^3,4^, posit that the brain represents the world with an internal generative model, used to predict future external (sensory input) and internal (physiological states) events. Model and cue-based expectations (prior beliefs) computationally combine with incoming sensory information (likelihoods) to form perceptions (posterior beliefs) ^5,6^. In line with this principle, predictions about upcoming stimuli have been shown to shape their perception across sensory modalities. For example, reported painfulness of noxious stimuli assimilates to expected values ^7–12^. Likewise, the perceived direction of randomly moving dots is biased towards expected directions ^5,13,14^. Such assimilation is beneficial, as it improves perception when contextual information is relevant. Similar Bayesian principles have been suggested to underlie interoception ^15–17^ and diverse aspects of cognition ^18,19^ including planning and decision-making (e.g., active inference ^20,21^). However, such models also have contrastive mechanisms, which drive learning. While assimilation of perceptions to predicted values is one way to resolve discrepancy between predictions and sensory input, another important way is to update the predictive model to match the input, i.e., to learn. Prediction error (PE)-driven updating is a crucial form of learning. However, which brain signals assimilate to expectations, perhaps forming a neural substrate for perception, and which encode PEs and thus drive learning, is a topic of active investigation.

Bayesian models of predictive processing make predictions about how the relative precision of cues and sensory stimuli affects perception and neural responses. Prior predictions and likelihoods (i.e., the expectation and the incoming sensory information) are weighted by their relative precision ^2,5,6,22–24^. Thus, Bayesian models predict an interaction between the expected value and its precision, such that more certain expectations (prior beliefs) should affect perception (posterior beliefs) more strongly. This hypothesis has been supported by studies on pain perception showing stronger placebo analgesia in individuals with more precise treatment expectations ^25,26^, and greater assimilation of perceived pain towards expectations with higher cue precision ^27,28^. Conversely, other studies have found that lower precision or predictability (higher uncertainty) leads to more pain, putatively because uncertainty is aversive in the context of potential harmful stimuli ^7,29^. Finally, some other studies did not replicate these effects and even found instead that more certain cues led to more pain ^30–34^. The contradicting findings regarding the effect of the cue certainty on pain perception ^35^ may stem from differences across experimental designs (e.g., how precision was manipulated, or the rating scale that was used), sensory modalities (e.g., uncertainty may be particularly aversive in the context of painful stimuli given the threat they pose), and / or participant populations (e.g., individual differences in the aversiveness of uncertainty, or participants’ beliefs about the information they are given and thus their generated expectations).

A promising paradigm, which we adopt here, uses cues with rich information about the distribution of predicted values, allowing within-person manipulation of precision and other distributional properties ^7,8,10,27,30^. This can substantially increase power and reduce confounds in testing manipulations of precision. The version that we use presents multiple-valued cues as ratings of multiple previous participants (Figure 1), providing an additional advantage in that it provides a plausible type of social observational learning that may be particularly impactful ^36–41^. Though such cues have been used successfully in multiple studies, an additional complexity is that manipulation of the cues’ precision–i.e., the variance of the presented cue values–influences other properties, including the maximum and minimum values. If people weight all cue values equally, this is ignorable, but participants may instead attend to the highest or lowest values, depending on the task and their predispositions (e.g., if some participants find pain to be threatening and are vigilant for high-pain cues).

**Figure 1.**
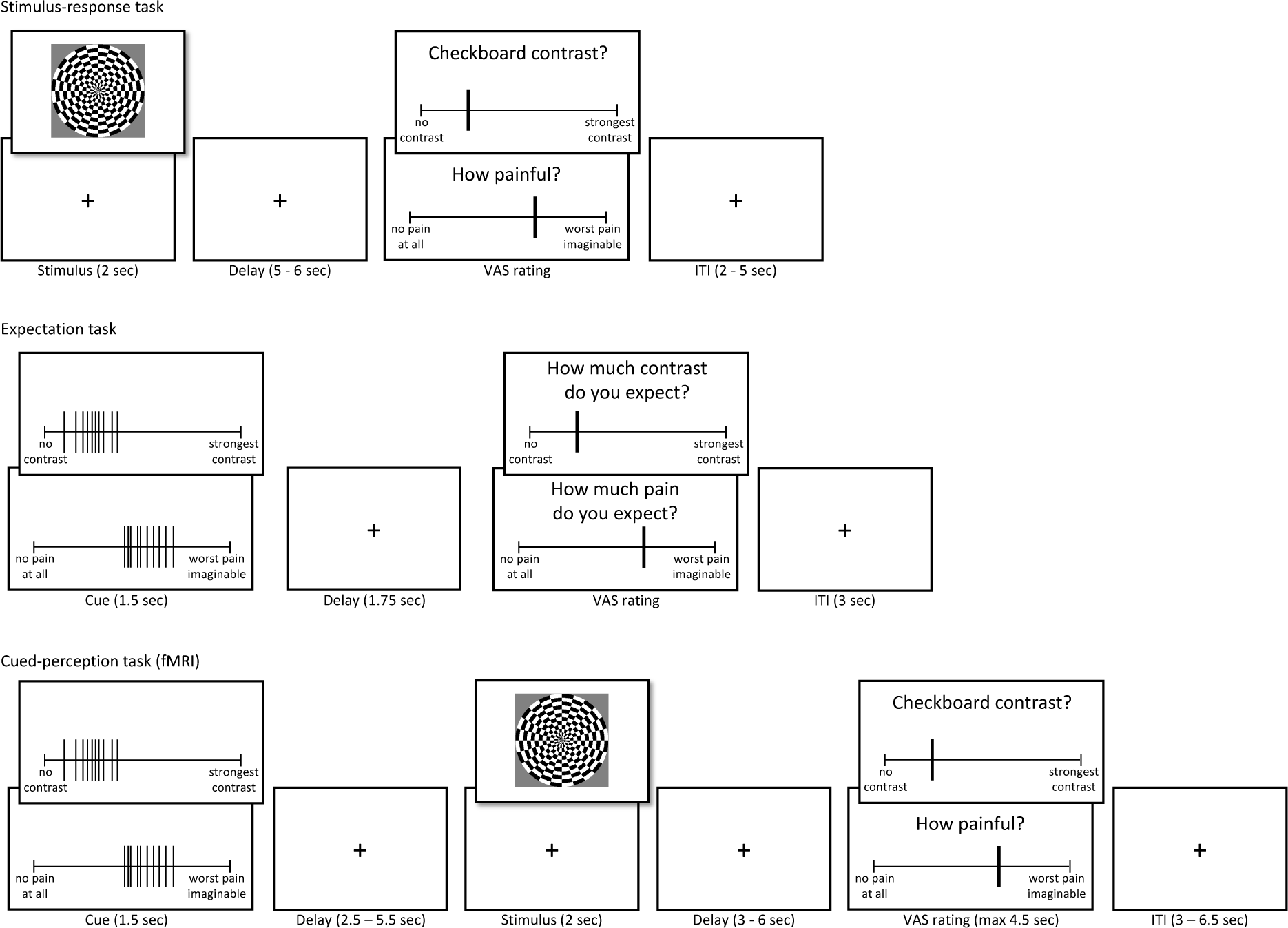
Experimental design. An illustration of the three experimental tasks: stimulus-response task (top panel), expectation task (middle panel), and cued-perception task (bottom panel; performed during an fMRI scan). When two rectangles are presented on top of each other (e.g., for the ratings in all tasks), the lower one illustrates pain trials and the upper one illustrates visual trials.

Such effects are not taken into consideration in most studies testing Bayesian accounts of perception, but there is evidence that they can be important. For instance, a previous study in visual perception has demonstrated overweighting of inliers (values closer to the mean) ^42^. Such effects may also be sensitive to the precise demands of the task. For example, when asked to determine which series of numbers was on average larger, people have been found to overweight outliers (extreme values), and specifically larger values ^43^. Furthermore, this kind of nonlinear weighting might depend on the sensory modality, and has not been studied in the context of pain perception. If attention to extreme values is related to vigilance for potential threat, people might weight them more strongly for pain than for other, safer modalities. This could, in principle, explain previous findings of higher pain ratings following less precise cues ^7^: If people are biased towards maximal pain values (“danger signals”), and higher variance entails larger maximal values, then this effect is expected.

Moreover, pain perception is considered more subjective and ambiguous compared to other types of perception, like perception of visual contrast. Thus, its likelihood might be less precise, and predictive cue effects might be increased compared to other modalities. If this is the case, it suggests that the nature of the predictive processes at play are at least to some degree modality-specific, and theories of perception must identify differences and commonalities across different types of sensory input ^23^.

Finally, it is still unclear how expectations, and their precision, influence brain processes underlying pain and visual perception. Some evidence suggest that influences of expectations can reach the earliest stages of sensory processing, for example in the spinal cord ^44–46^, cortical nociceptive pain processing regions ^9,47^, and primary visual cortex ^14,48^. Other studies show that expectations modulate higher-level affective and cognitive processes ^49,50^. However, others have questioned how strongly context-based influences affect perception ^51^. Moreover, we have recently shown that placebo treatment (which is thought to operate via expectation modulation, among other effects ^52–54^) induces analgesia via modulation of affective and cognitive processes, rather than nociceptive pain processing ^55^. If expectation effects mostly operate via high-level affective and cognitive processes, they should be reduced for basic perception that is not subject to affective evaluation, like brightness or visual contrast. Alternatively, these processes could be mediated by different mechanisms across modalities, and thus subject to different behavioral influences and related to different patterns of neural activity.

Here, we test how expectations are generated from distribution cues with varying levels of mean, variance, and skewness (and thus the presence and direction of outliers), and how they affect subsequent perception of visual and painful stimuli (for the study design and illustration of the cues see Figure 1 and Methods). The cues were not externally reinforced and were not predictive of subsequent stimuli. By comparing pain and visual perception, we identify common and distinct mechanisms for expectation formation and modulation of perception for stimuli that are potentially harmful and more ambiguous (painful) and those that are non-harmful and less ambiguous (visual). Using fMRI and a priori neuromarkers and individual regions of interest (ROIs), we test whether and how neural responses during pain and visual perception are affected by the different properties of the cue and by the cue-based expectations.

## Results

In the first task, participants rated the painfulness of thermal stimuli and the visual contrast of flickering checkerboards with different intensities (Figure 1). Data from this task verified that participants’ ratings were higher for higher stimulus intensity levels in both modalities (linear mixed-effects model: pain: *β* = 0.633, *SE* = 0.028, *t*_(44)_ = 22.43, *p* < .001; vision: *β* = 0.868, *SE* = 0.018, *t*_(44)_ = 47.59, *p* < .001).

### Expectation generation from distribution cues

In the expectation task, participants were presented with the distribution cues, and rated their expectations regarding noxious and visual stimuli following each cue (stimuli were not delivered during this task).

#### Effects of cue mean on expectations

Expectation ratings significantly assimilated to the cues’ mean (linear mixed-effects model: *β* = 0.907, *SE* = 0.015, *t*_(50)_ = 59.14, *p* < .001; Figure 2A). This effect did not interact with modality (*β* = -0.006, *SE* = 0.005, *t*_(16000)_ = -1.26, *p* = .210), and was found for each modality separately (pain: *β* = 0.901, *SE* = 0.015, *t*_(57.7)_ = 62.72, *p* < .001; vision: *β* = 0.913, *SE* = 0.021, *t*_(51.3)_ = 43.25, *p* < .001).

**Figure 2.**
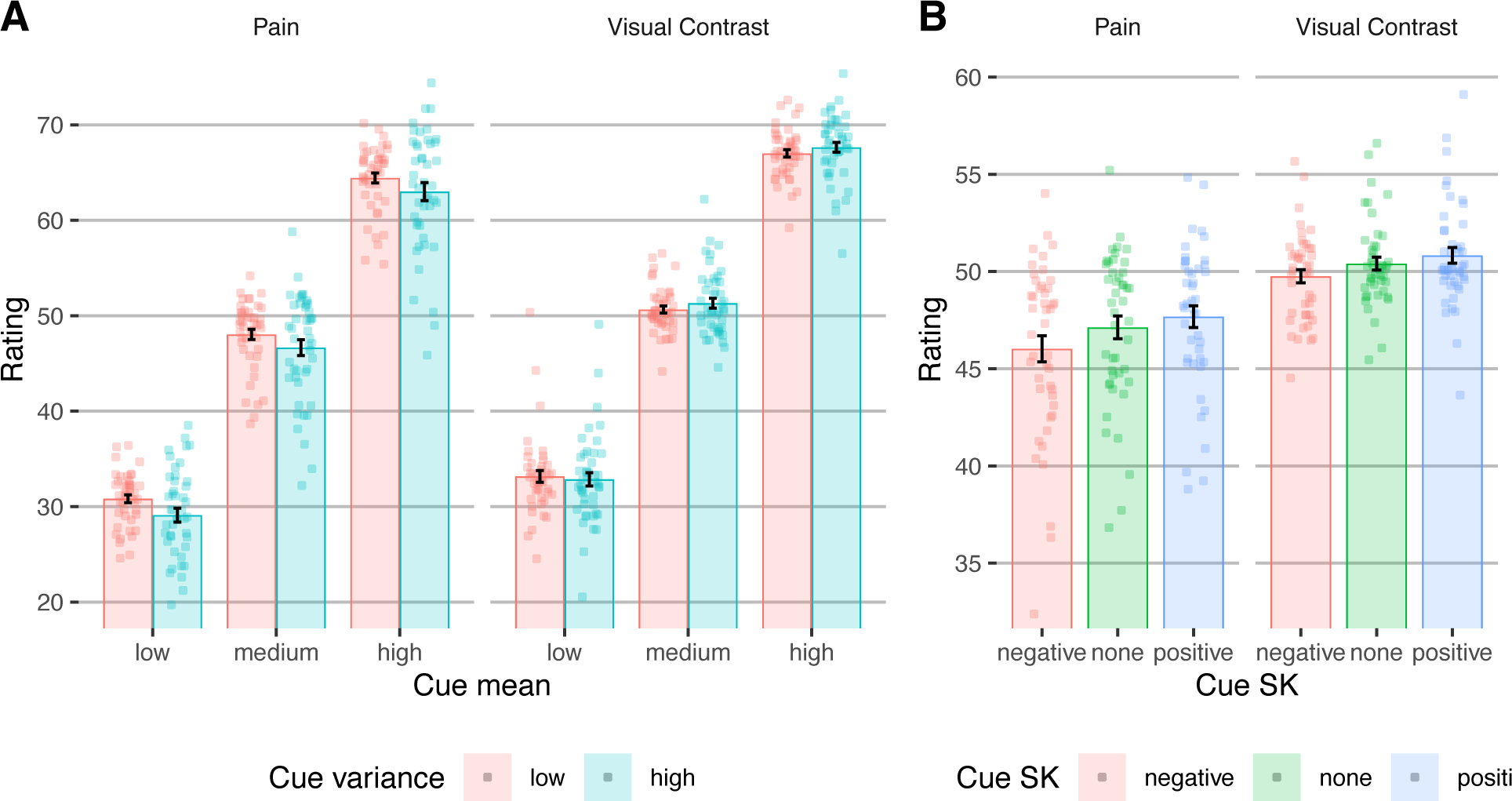
Behavioral results - expectation task. (**A**) Participants’ expectation ratings as a function of the mean (five cue mean levels were collapsed into three for visualization) and variance of the cue’s values, in pain and vision trials. (**B**) Participants’ expectation ratings as a function of the skewness of the cue’s values, in pain and vision trials. In both panels, bars represent averages across participants, error bars represent the standard error of the mean across participants, and points represent single participants.

#### Effects of cue variance on expectations

Previous studies have mostly focused on the effect of cues’ variance on perception rather than expectations, as the variance should influence the precision (certainty) of the expectations, not their value. Here, the cue variance did not significantly affect expectation ratings across both modalities (*β* = -0.021, *SE* = 0.010, *t*_(59.83)_ = - 1.994, *p* = .051; Figure 2A), but there was a significant cue mean x modality interaction (*β* = - 0.024, *SE* = 0.005, *t*_(16000)_ = -5.01, *p* < .001): Expectation ratings were significantly higher for lower cue variance in pain trials (*β* = -0.045, *SE* = 0.014, *t*_(58.1)_ = -3.16, *p* = .002), but not in vision trials (*β* = 0.004, *SE* = 0.013, *t*_(66)_ = 0.25, *p* = .802). In other words, participants expected more painful stimuli after more certain cues, but this was not true for noxious stimuli. This finding is inconsistent with both the Bayesian predictive processing account and the alternative account of uncertainty aversiveness. We further tested the interaction between the effects of cue mean and variance on subsequent ratings, which based on Bayesian predictive processing accounts is expected to be significant for stimulus, and not expectation, ratings. Indeed, the cue mean x variance interaction was not significant in either modality (pain: *β* = -0.0002, *SE* = 0.006, *t*_(7956)_ = -0.026, *p* = .979; vision: *β* = 0.014, *SE* = 0.007, *t*_(7956)_ = 1.93, *p* = .053).

#### Effects of cue skewness on expectations

We tested for the first time how the skewness of the cue distribution affects expectations. We reasoned that previous studies that have manipulated the cue variance also altered extreme values, and that people might be biased towards such values, particularly in the context of threat (i.e., on pain trials). Here, extreme values indeed biased participants’ expectations, which were higher for positively skewed compared to symmetric cues (*β* = 0.028, *SE* = 0.007, *t*_(16000)_ = 4.11, *p* < .001; Figure 2B) and lower for negatively skewed compared to symmetric cues (*β* = -0.050, *SE* = 0.007, *t*_(16000)_ = -7.38, *p* < .001). These effects were also significant in each modality separately (pain, negative vs. symmetric: *β* = -0.064, *SE* = 0.009, *t*_(7956)_ = -7.13, *p* < .001; pain, positive vs. symmetric: *β* = 0.032, *SE* = 0.009, *t*_(7956)_ = 3.59, *p* < .001; vision, negative vs. symmetric: *β* = -0.037, *SE* = 0.010, *t*_(7956)_ = -3.77, *p* < .001; vision, positive vs. symmetric: *β* = 0.024, *SE* = 0.010, *t*_(7956)_ = 2.45, *p* = .014). These findings are consistent with over-weighting of extreme values when constructing expectations.

In addition, expectation ratings were significantly higher in vision compared to pain trials (Vision: *M* = 50.33, *SE* = 0.319; Pain: *M* = 46.945, *SE* = 0.586; *β* = -0.094, *SE* = 0.020, *t*_(47.8)_ = -4.82, *p* < .001). This result suggests that participants might have been biased by lower values particularly in the context of pain, or viewed themselves as less sensitive to pain compared to other participants. Finally, there was a significant cue variance x cue skewness interaction (*β* = 0.021, *SE* = 0.007, *t*_(16000)_ = 3.064, *p* = .002), indicating that lower extreme values affected expectations more when there was overall stronger agreement (less variance) among the cue values.

Overall, the behavioral results of the expectation task suggest that extreme values bias expectations, and that participants were particularly attentive to lower extreme values, mostly in the context of pain.

#### Computational model: weighting of cue values

To more directly test whether participants weight all cue values equally or overweight particular values (e.g., inliers vs. outliers, or smaller vs. larger values), we developed a computational model of expectation generation. The model assumes that in each cue, each value is weighted based on its relative location in the distribution of the 10 values (Figure 3A). Each value’s weight is based on a combination of a power term modeling the weighting of inliers vs. outliers with the free parameter *k*, and a logistic term modeling the weighting of values that are smaller vs. larger than the mean with the free parameter *b* (see Methods for a detailed description). The model was largely inspired by the model of Spitzer et al. in the context of numeric estimation ^43^.

**Figure 3.**
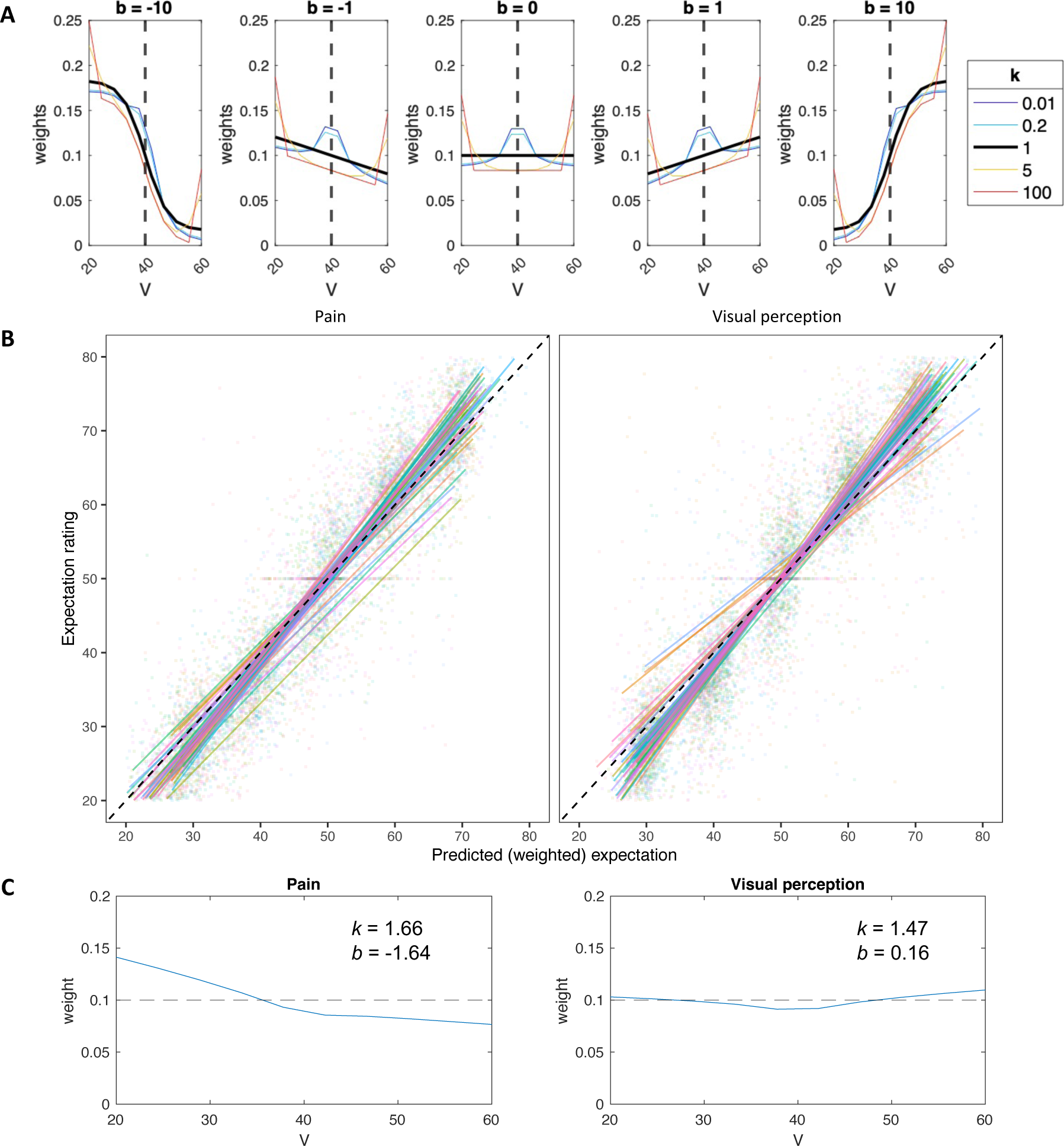
Computational model of expectation generation. (**A**) Simulations of the expectation model: Mapping of cue values, *V* (10 per cue), to weights for expectation computation, based on the two free parameters of the model: *k* and *b*. When *k* = 1 (black line), inliers and outliers are equally weighted. When *k* < 1 (cold colors), inliers are over-weighted, and when *k* > 1 (hot colors), outliers are over-weighted. When *b* < 0 (left panels), values below the mean are over-weighted, when *b* > 0 (right panels) values above the mean are over-weighted, and when *b* = 0 (middle panel), values are equally weighted. The dashed line represents the cue mean. (**B**) Correlations between the observed and predicted (based on the computational model) expectation ratings were very high across participants. Each line represents a single participant, and each dot represents a single trial. Data from different participants are presented in different colors. (**C**) The weight function for each modality, based on the median *k* and *b* values across the group. The dashed line represents equal rating of all cue values (*V*). Note that panels A and C are based on a symmetric cue consisting of equally distributed values between 20 and 60.

We fit the model to the expectation data to estimate the two free parameters, *k* and *b* for each modality and participant. The expectation model fit the data well overall, with an average correlation between predicted and empirical expectation ratings across participants of Pearson’s *r* = 0.945 (*SD* = 0.032) for pain and *r* = 0.928 (*SD* = 0.074) for vision, and an average root mean squared error (RMSE) of 5.904 (*SD* = 1.668) for pain and 6.415 (*SD* = 2.545) for vision (on a rating scale of 0-100). This indicates that participants tracked variation in cues accurately, and that the model predicted their responses well and performed generally better in pain than in vision (Figure 3B).

The free parameter *k* measures over-weighting of inliers (*k* > 1) or over-weighting of outliers (*k* < 1), while *b* measures over-weighting of values that are smaller (*b* < 0) or larger (*b* > 0) than the mean value. We found that participants significantly over-weighted outliers in both modalities (i.e., *k* > 1; Wilcoxon signed rank test; Pain: median *k* = 1.66, *p* = .019; Vision: median *k* = 1.47, *p* = .008), and also smaller values specifically in pain and not in vision (i.e., *b* < 0; Wilcoxon signed rank test; Pain: median *b* = -1.64, *p* < .001; Vision: median *b* = 0.16, *p* = .446). Furthermore, *k* values, but not *b* values, were correlated between the two modalities across participants (Spearman correlation; *k*: *ρ* = 0.45, *p* = .002; *b*: *ρ* = 0.16, *p* = .304).

Finally, we tested whether the estimated *k* and *b* values correlated with state / trait scores based on questionnaires completed by the participants prior to the cued-perception task. For example, people with higher fear of pain, pain catastrophizing, or anxiety may focus more on cues predicting higher expected pain. However, all correlations between the available measures in our sample (Fear of Pain score ^56^, Pain Catastrophizing score ^57^, and State-Trait Anxiety Inventory score ^58^) and optimized *k* or *b* for each modality across participants, were not significant (Fear of Pain: *k* pain *r* = 0.16, *p* = .284, *k* vision *r* = 0.04, *p* = .800, *b* pain *r* = 0.22, *p* = .140, *b* vision *r* = 0.03, *p* = .825; Pain Catastrophizing: *k* pain *r* = -0.17, *p* = .267, *k* vision *r* = 0.09, *p* = .553, *b* pain *r* = 0.08, *p* = .582, *b* vision *r* = 0.06, *p* = .705; State Anxiety: *k* pain *r* = 0.15, *p* = .32, *k* vision *r* = 0.13, *p* = .402, *b* pain *r* = 0.05, *p* = .734, *b* vision *r* = -0.02, *p* = .905).

Taken together, the behavioral and computational results of the expectation task suggest that while expectations strongly assimilate towards the mean cue value, participants overweight extreme cue values in both modalities, and demonstrate an optimism bias specifically in pain, relying more on low than high extreme values. Such effects were not considered in most previous studies, and could potentially explain the inconsistent results with regard to the effect of expectation variance on pain perception, if extreme values were unintentionally manipulated along with the uncertainty.

### The effects of cue-based expectations on perception

In the cued-perception task (Figure 1), participants viewed cues followed by painful heat or flickering checkerboards. As expected, participants rated stimuli with higher intensity levels (higher temperature or visual contrast) as more intense (*β* = 0.322, *SE* = 0.016, *t*_(5856)_ = 20.46, *p* < .001). This effect was also significant for each modality separately (pain: *β* = 0.323, *SE* = 0.027, *t*_(129.6)_ = 11.838, *p* < .001; visual perception: *β* = 0.333, *SE* = 0.032, *t*_(88.8)_ = 10.552, *p* < .001).

#### Effects of cue mean on perception

Participants’ stimuli ratings assimilated to the cue mean (*β* = 0.253, *SE* = 0.033, *t*_(61.5)_ = 7.693, *p* < .001; Figure 4A), replicating previous studies. This effect was significant for both pain (*β* = 0.230, *SE* = 0.036, *t*_(81.44)_ = 6.443, *p* < .001) and visual perception (*β* = 0.289, *SE* = 0.039, *t*_(69.6)_ = 7.503, *p* < .001), and cue mean effects on pain and visual perception were correlated across participants (Pearson’s r = 0.82, *t*_41_ = 9.174, 95% CI = [0.689, 0.899], *p* < .001), indicating that participants who were affected by the cues were similarly affected in both modalities.

**Figure 4.**
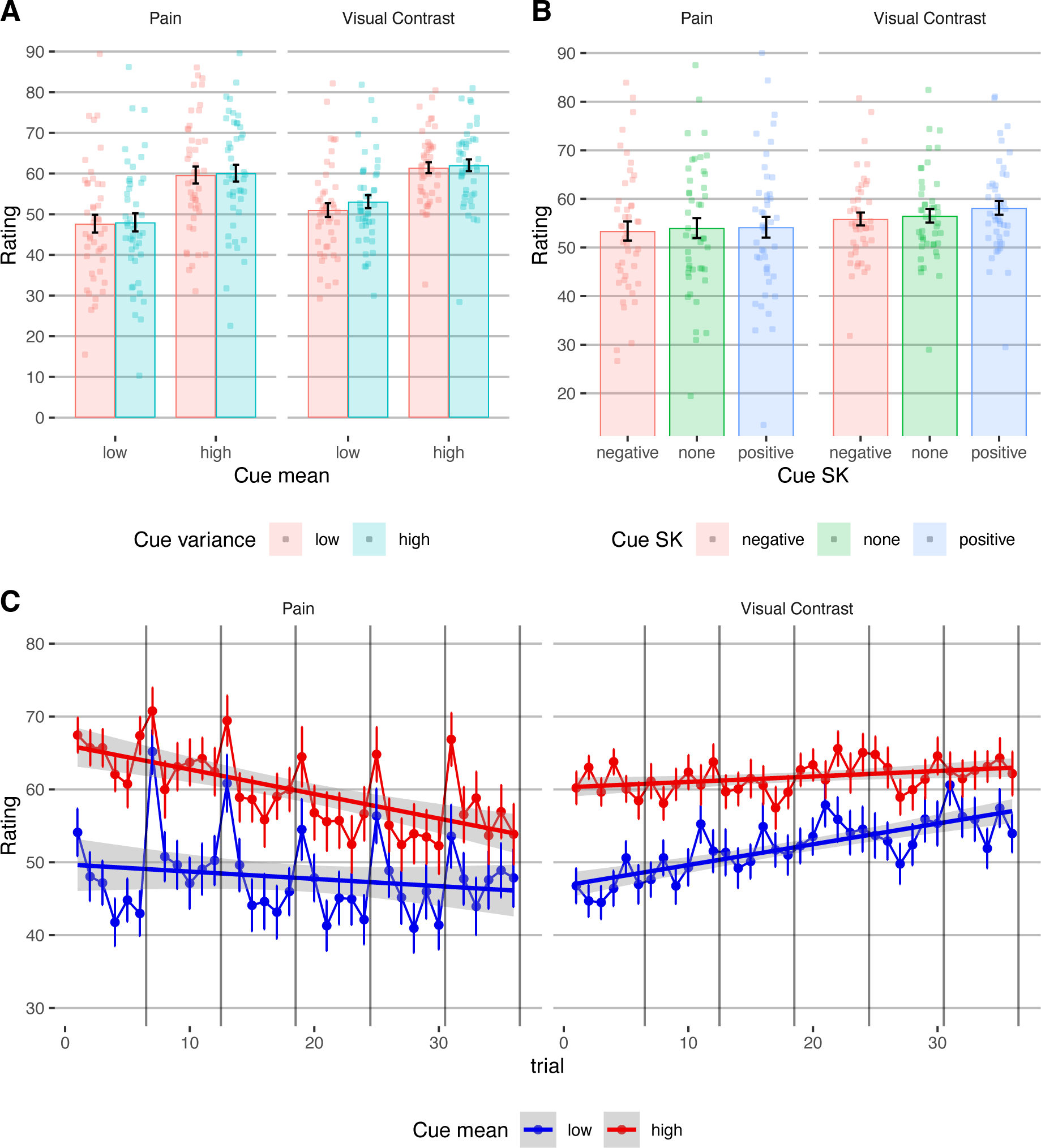
Behavioral results, cued-perception task. (**A**) Participants’ perception ratings as a function of the mean (x axis) and variance (color) of the cue’s values, in pain and visual trials (column). (**B**) Participants’ perception ratings as a function of the skewness (x axis and color) of the cue’s values, in pain and visual trials (columns). In panels A and B, bars represent averages across participants, error bars represent the standard error of the mean across participants, and points represent single participants. (**C**) The effect of the cue mean (color) on pain and visual contrast ratings over time, in pain and visual perception (columns). Error bars represent standard error of the mean across participants. The lines represent the linear fit, and the gray shading represents 95% CIs of the linear fit. Vertical major grids separate between different runs (note that a new skin site was used for each run, and thus the increase in the averaged pain rating for the first trial of each run stems from site-nonspecific sensitization and site-specific habituation^59^).

#### Effects of cue variance on perception

Participants rated stimuli with higher cue variance as more intense (*β* = 0.037, *SE* = 0.016, *t*_(5857)_ = 2.320, *p* = .020). This finding is consistent with Yoshida et al., ^7^, who suggested that uncertainty is aversive in the context of pain and thus lead to higher pain ratings, although a direct replication did not find this effect ^30^. The modality x cue variance interaction was not significant (*β* = -0.137, *SE* = 0.016, *t*_(5858)_ = -0.869, *p* = .385). However, when tested separately, it was only significant for visual (*β* = 0.060, *SE* = 0.021, *t*_(2877)_ = 2.837, *p* = .005) and not for pain stimuli (*β* = 0.021, *SE* = 0.022, *t*_(2846)_ = 0.928, *p* = .354), which does not support the hypothesis that uncertainty is aversive particularly in the context of harmful stimuli.

As described above, Bayesian predictive processing accounts predict that the effect of the cue mean (expectation value) on perception should be smaller with higher cue variance (lower expectation precision). In line with this prediction, across modalities, the effect of the cue mean was smaller when the cue variance was higher (cue mean *x* variance interaction: *β* = -0.044, *SE* = 0.016, *t*_(5859)_ = -2.828, *p* = .005). This effect did not interact with modality (*β* = 0.003, *SE* = 0.016, *t*_(5857)_ = 0.212, *p* = .832), but when tested separately, it was only significant for visual stimuli (*β* = -0.056, *SE* = 0.021, *t*_(2878)_ = -2.661, *p* = .008) and not for pain (*β* = -0.036, *SE* = 0.022, *t*_(2850)_ = -1.594, *p* = .111).

#### Effects of cue skewness on perception

Ratings were higher when they were preceded by positively skewed compared to symmetric cues (*β* = 0.046, *SE* = 0.022, *t*_(5857)_ = 2.052, *p* = .040; Figure 4B), but were not lower following negatively skewed compared to symmetric cues (*β* = - 0.034, *SE* = 0.022, *t*_(5857)_ = -1.547, *p* = .122). The modality x skewness interactions were not significant (negative vs. symmetric: *β* = -0.002, *SE* = 0.022, *t*_(5857)_ = -0.104, *p* = .917; positive vs. symmetric: *β* = -0.032, *SE* = 0.022, *t*_(5857)_ = -1.421, *p* = .155). However, when tested separately, the skewness only affected visual perception (higher ratings after positively skewed vs. symmetric cues; *β* = 0.092, *SE* = 0.030, *t*_(2878)_ = 3.086, *p* = .002; no significant effect for negatively skewed vs. symmetric cues, *β* = -0.038, *SE* = 0.030, *t*_(2876)_ = -1.276, *p* = .202) and not pain (negative vs. symmetric: *β* = -0.033, *SE* = 0.032, *t*_(2845)_ = -1.038, *p* = .299; positive vs. symmetric: *β* = 0.012, *SE* = 0.032, *t*_(2845)_ = 0.377, *p* = .706).

Overall, ratings were higher following cues with higher mean, higher variance, or positive skewness, suggesting that extreme positive values carry more influence on perception. Surprisingly, however, effects of the cue precision and extreme values on perception were mostly found in visual rather than pain perception, inconsistent with the idea that uncertainty enhances threat. In line with Bayesian predictive processing accounts, the cue mean influenced perception more when the cues were more precise. However, predictive processing accounts do not explain why the ratings were also higher following higher uncertainty, and why the cue precision was not a significant factor in pain perception. Finally, computational modeling indicated that perception assimilates towards expected values more in pain compared to visual perception, in line with the hypothesis that pain perception is more ambiguous and thus more sensitive to contextual information (see Supplementary Results - Computational Modeling of Perception).

### Does the effect of cue-based expectation persist?

The interaction between the trial number and the cue mean effect was not significant (Figure 4C; both modalities: *β* = -0.009, *SE* = 0.011, *t*_(5918)_ = -0.816, *p* = .414; pain: *β* = -0.002, *SE* = 0.015, *t*_(2903)_ = -0.115, *p* = .908; visual perception: *β* = -0.010, *SE* = 0.015, *t*_(2948)_ = -0.669, *p* = .503), and the cue mean effect was significant on the last trial (*t_81_* = -3.18, mean difference 95% CI = [-13.427, -3.101], *p* < .001), demonstrating a persistent cue effect. Computational modeling strengthened the conclusion that most participants did not learn to ignore the cues during the task (see Supplementary Results - Computational Modeling of Perception), in line with previous studies ^9,60–62^.

### Cue and stimulus intensity effects on neural processing

#### Effects on pain neuromarkers

We focused on two *a priori* neuromarkers for pain (Figure 5A): (1) The Neurologic Pain Signature (NPS ^63^), which is sensitive and specific to nociceptive pain across studies, tracks the intensity of nociceptive input, and predicts pain ratings with very large effect sizes in >50 published study cohorts ^49,64^. (2) The Stimulus Intensity Independent Pain Signature (SIIPS ^65^), which captures higher-level, endogenous influences on pain construction independent of stimulus intensity and the NPS score. In most previous studies, the NPS was not modulated by expectations or other contextual effects ^55,66^, although some previous studies have found modulations with some types of interventions ^62,67^, sometimes with very small effect size ^49^.

**Figure 5.**
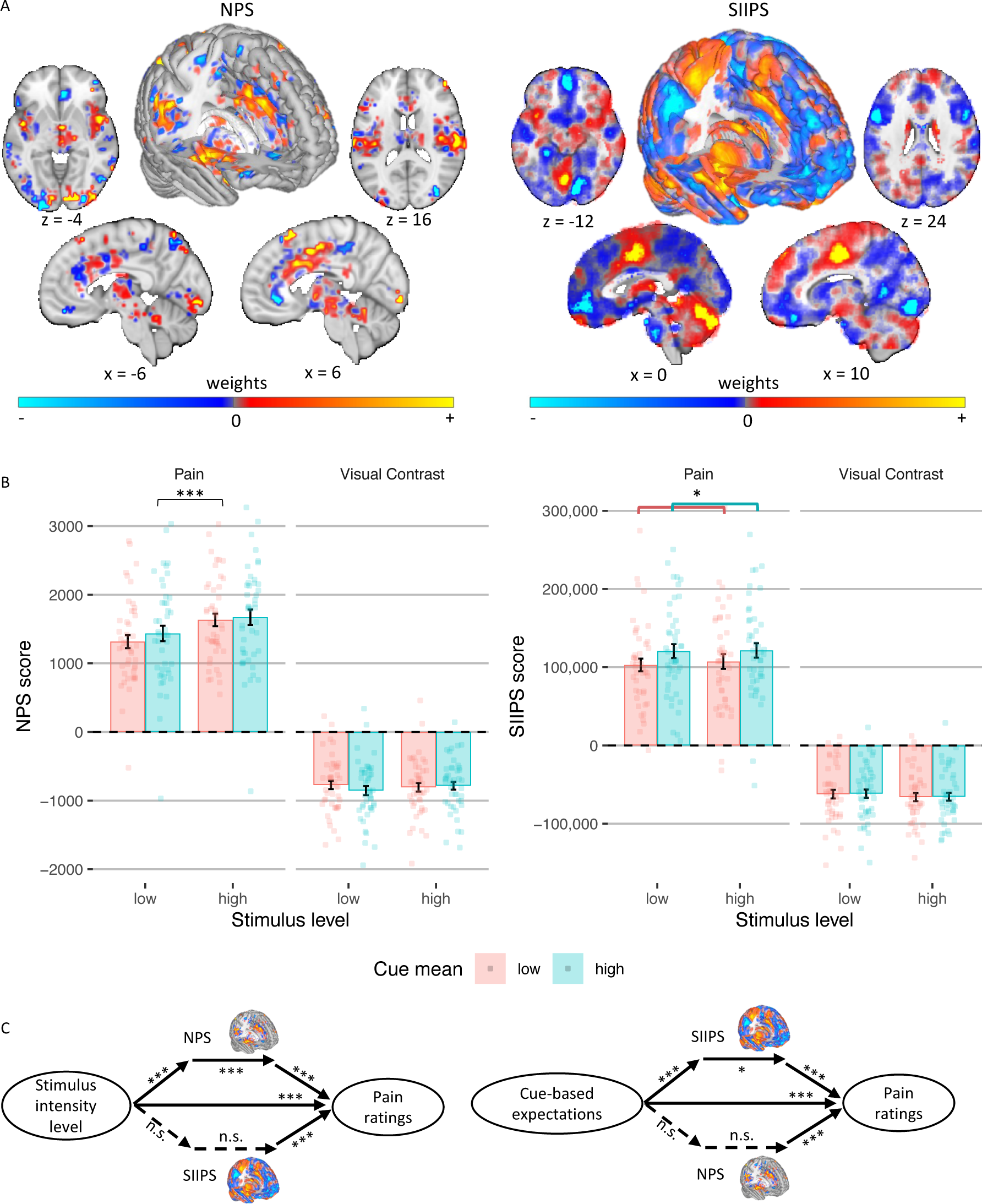
Neuromarker results. (**A**) The NPS and SIIPS neuromarkers. (**B**) NPS (left) and SIIPS (right) score as a function of the stimulus intensity (x axis), cue mean (color) and modality (column). Bars represent averages across participants, error bars represent the standard error of the mean across participants, and points represent single participants. (**C**) Multilevel mediation analysis with pain neuromarkers. Solid lines represent significant effects and dashed lines represent non-significant effects. Asterisks represent the level of significance (* *p* < .05, *** *p* < .001).

Conversely, the SIIPS was found to be affected by expectations and related psychological manipulations such as placebo treatment ^55^, perceived control and conditioned cues ^66^. Notably, a study using social distribution cues like the ones we have used here found no effects of the cues on the NPS and SIIPS scores ^10^.

We computed a dot product based score, the ‘pattern response’, for each trial, and included the scores for each neuromarker in mixed-effects models identical to the ones used for the behavioral pain ratings. The NPS score was significantly higher for high compared to low intensity painful stimuli (*β* = 0.143, *SE* = 0.027, *t*_(2895.21)_ = 5.23, *p* < .001), replicating previous studies. However, it was not significantly affected by the cue mean (*β* = 0.022, *SE* = 0.030, *t*_(202.09)_ = 0.74, *p* = .462), variance (*β* = 0.006, *SE* = 0.027, *t*_(2895.26)_ = 0.24, *p* = .812), or skewness (positive vs. symmetric: *β* = 0.017, *SE* = 0.038, *t*_(2895.6)_ = 0.45, *p* = .653; symmetric vs. negative: *β* = 0.047, *SE* = 0.039, *t*_(2894.92)_ = 1.21, *p* = .225). The cue mean x cue variance interaction was also not significant (*β* = 0.006, *SE* = 0.027, *t*_(2904.36)_ = 0.21, *p* = .836). Thus, the NPS response depends on the noxious input, and is not modulated by the cues (Figure 5B).

Conversely, the SIIPS score assimilated towards the cue mean (*β* = 0.066, *SE* = 0.028, *t*_(2936.51)_ = 2.33, *p* = .020; (Figure 5B)), but was not significantly affected by the stimulus intensity (*β* = - 0.017, *SE* = 0.028, *t*_(2936.19)_ = -0.59, *p* = .552), cue variance (*β* = 0.016, *SE* = 0.028, *t*_(2937.62)_ = 0.55, *p* = .586), or cue skewness (positive vs. symmetric: *β* = -0.012, *SE* = 0.040, *t*_(2937.06)_ = -0.31, *p* = .757; symmetric vs. negative: *β* = -0.011, *SE* = 0.040, *t*_(2936.81)_ = -0.27, *p* = .785). The cue mean x cue variance interaction was not significant (*β* = -0.007, *SE* = 0.028, *t*_(2936.44)_ = -0.24, *p* = .814). Importantly, the NPS and SIIPS are pain neuromarkers, and should not respond to neutral visual stimuli. Indeed, when testing the NPS and SIIPS scores during perception of visual stimuli, there were no significant effects of stimulus intensity or cues (all *p*s ≥ .094; Supplementary Table 2).

#### Multilevel mediation analysis

We used mediation analysis to test whether the NPS and / or SIIPS formally mediate the effect of the cue-based expectations and/or stimulus intensity on trial-by-trial pain reports (Figure 5C). Trial-by-trial expectancy scores were based on the computational model (see “Computational model: weighting of cue values” above). Expectancy models controlled for stimulus intensity as a covariate, and vice versa.

The NPS partially mediated the effect of stimulus intensity level on pain ratings (*path ab* stimulus intensity level → NPS score → pain rating: *β* = 0.02, *SE* = 0.00, *z* = 3.68, *p* < .001), controlling for cue-based expectations. Higher stimulus intensities led to higher NPS scores (*path a*, *β* = 0.11, *SE* = 0.01, *z* = 4.00, *p* < .001), and higher NPS scores predicted greater trial-by-trial pain (*path b, β* = 0.28, *SE* = 0.02, *z* = 3.85, *p* < .001). However, the NPS did not mediate the effect of the cue-based expectations on pain ratings (*path ab* cue-based expectation → NPS score → pain rating: *β* = 0.00, *SE* = 0.00, *z* = 1.42, *p* = .154), controlling for stimulus intensity level, since higher cue-based expectations did not lead to higher NPS scores (*path a*, *β* = 0.02, *SE* = 0.02, *z* = 1.24, *p* = .216).

Conversely, the SIIPS partially mediated the effect of the cue-based expectation on pain ratings (*path ab* cue-based expectation → SIIPS score → pain rating: *β* = 0.003, *SE* = 0.001, *z* = 2.45, *p* = .014), controlling for stimulus intensity level. Higher cue-based expectations led to higher SIIPS scores (*path a, β* = 0.06, *SE* = 0.01, *z* = 3.88, *p* < .001), and higher SIIPS scores predicted greater trial-by-trial pain (*path b* SIIPS score → pain rating: *β* = 0.15, *SE* = 0.03, *z* = 3.82, *p* < .001). The SIIPS did not mediate the effect of the stimulus intensity level on pain ratings (*path ab* stimulus intensity level → SIIPS score → pain rating: *β* = 0.00, *SE* = 0.00, *z* = 1.37, *p* = .171), controlling for cue-based expectations, since stimulus intensity was not associated with SIIPS scores (*path a*, *β* = 0.02, *SE* = 0.01, *z* = 1.23, *p* = .219).

As expected, neither neuromarker mediated the effect of the stimulus intensity level or the effect of the cue-based expectation on the contrast rating of visual stimuli. Overall, the NPS score was only affected by the heat intensity and formally partially mediated the effect of the stimulus intensity on pain ratings, while the SIIPS score assimilated to the cue mean and partially mediated the effect of the cue-based expectations on pain ratings. Other properties of the cue, including the cue variance, skewness, and the interaction between the cue mean and cue variance, did not affect either neuromarker. These results indicate that the cues did not affect pain via changes in early nociceptive pain processing, but rather via systems associated with endogenous contributions to pain perception ^55^.

#### Effects on ROIs related to early neural perceptual processing

To complement neuromarker analyses, which capture activity in distributed systems, we tested whether the cues affected activity in several a priori ROIs related to pain and visual perception (Figure 6; for all regions see Supplementary Table 1). We first focused on regions associated with early perceptual representations (for full statistics see Supplementary Table 3 and Supplementary Table 4). For nociception, we focused on the spinothalamic tract, including ventral posterior (VPL/VPM) thalamus and dorsal posterior insula (dpIns). Activity in these regions was larger for more intense heat stimuli (all *p*s ≤ .003), but was not affected by the cue mean, cue variance, cue skewness, or cue mean x cue variance interaction (all *p*s ≥ 0.141). For early visual perception, we focused on the lateral geniculate nucleus (LGN) and primary and secondary visual cortex (V1 and V2). Activity was higher for higher visual contrast in the right V1 (*p* = .041) and left V2 (*p* = .028), but not in the LGN (right *p* = .438; left *p* = .332), left V1 (*p* = .067) and right V2 (*p* = .058). Like early nociceptive pain perception, early visual perception was not affected by the cues (negative vs. symmetric cues in right V2 *p* = .072 and left V2 *p* = .056; all other *p*s > 0.153). Thus, in sum, most of these areas showed stimulus intensity effects, but none were influenced by the cues.

**Figure 6.**
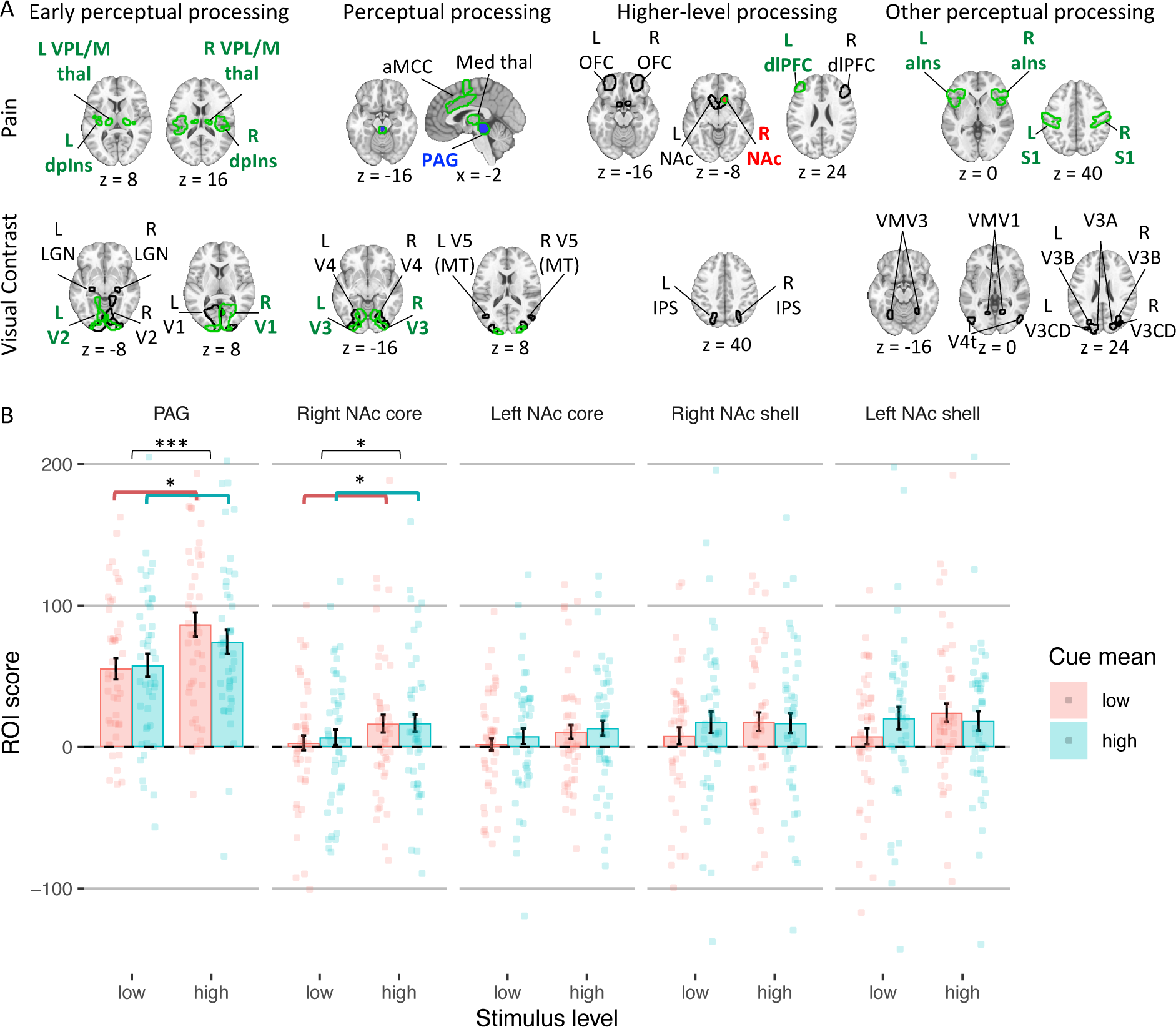
ROI results. (**A**) ROIs are shown with a contour for each modality (rows) and processing stage (columns). Regions with higher activity for more intense stimuli are presented with a green contour. Regions with a significant cue mean effect are presented in red (higher activity for higher cue mean) or blue (higher activity for lower cue mean). (**B**) The ROI score in the two regions with significant cue mean effect during pain perception (including other parts of the NAc) as a function of the stimulus intensity level (x axis) and cue mean (color). Bars represent averages across participants, error bars represent standard error of the mean across participants, and points represent single participants. Asterisks represent the level of significance (* *p* < .05, *** *p* < .001). Abbreviations: aIns = anterior insula; aMCC = anterior midcingulate cortex; dlPFC = dorsolateral prefrontal cortex; dpIns = dorsal posterior insula; L = left; Med = medial; NAc = nucleus accumbens; OFC = orbitofrontal cortex; PAG = periaqueductal gray; R = right; thal = thalamus.

#### Effects on ROIs related to neural perceptual processing

We then tested cue effects on regions associated with higher-level perceptual processing and affect (Figure 6; see Supplementary Tables 5 and 6 for full statistics). For pain, these included the periaqueductal gray (PAG), anterior midcingulate cortex (aMCC), and medial thalamus. For visual perception they included V3, V4 and V5 (MT). Activity in all pain processing regions was again higher for higher noxious stimulus intensity (all *p*s ≤ .001), but only the PAG was sensitive to predictive cues: PAG activity was higher following low-vs. high-mean cues (*p* = .031), consistent with aversive PE-like responses found in the PAG in previous studies ^68,69^. In the visual perception task, activity in visual regions was only higher for higher visual contrast in V3 (right *p* = .046, left *p* = .017; all other *p*s ≥ .193). While the cue mean, cue variance, and the interaction between them was not significant for any of these visual regions (all *p*s ≥ .106), the cues’ skewness did affect activity: Activity was lower following negatively skewed compared to symmetric cues in bilateral V3 (right, *p* = .009; left, *p* = .032), bilateral V4 (right, *p* = .010; left, *p* = .045), and left V5 (*p* = .027). Activity was not different following positively skewed compared to symmetric cues in all regions (all *p*s ≥ .111). Overall, the results validate the PAG’s role in PEs, but provide no evidence for assimilation to cue value or encoding of cue precision in pain, and only weak evidence for assimilation to extreme low-value cues in visual areas.

#### Effects on ROIs related to higher-level processing

Next, we tested regions associated with higher-level affective and cognitive processing of noxious and visual stimuli, including the nucleus accumbens (NAc; shell-like and core-like parts), dorsolateral prefrontal cortex (dlPFC) and mid-lateral orbitofrontal cortex (OFC) for pain, and the intraparietal sulcus (IPS) for visual perception (Figure 6; see Supplementary Tables 7 and 8 for full statistics). Activity was higher for more intense heat stimuli in the right NAc (core-like part; *p* = .017) and left dlPFC (*p* = .001). The cues only affected activity in the NAc (core-like part; all other *p*s ≥ .070): Activity in the right NAc was higher following cues with a higher mean (*p* = .018) and lower variance (i.e., more certain; *p* = .011), and activity in the left NAc was higher for low-variance (more precise) cues (*p* = .001) and showed a cue mean *x* cue variance interaction (*p* = .015), such that cue mean effects were smaller for more precise cues. These findings were not in line with previous predictive coding findings _25,70_ (see Discussion). In visual perception, activity in the IPS was not affected by the stimulus intensity or the cues (cue mean effect: *p* = .066 in the left and *p* = .074 in the right IPS; cue mean x cue variance interaction in the right IPS *p* = .064; all other *p*s ≥ .216).

#### Effects on other perceptual processing regions

Finally, we performed an exploratory analysis testing a larger set of regions, including anterior insula (aIns) and primary somatosensory cortex (S1, mostly the hand area) for pain processing, and V3A, V3B, V3CD, V4t, VMV1, VMV2, and VMV3 for visual processing (Figure 6; see Supplementary Tables 9 and 10 for full statistics). Activity during pain perception was higher for higher stimulus intensity in the aIns (right *p* = .007; left *p* = .001) and S1 (both *p* < .001), but the cues did not affect their activity (negatively skewed vs. symmetric cues in the right aIns *p* = .075, all other *p*s ≥ 0.225). Interestingly, activity was higher (less negative) with more intense heat in several visual regions, possibly because noxious heat stimuli produce diffuse effects on neuromodulatory systems, but none showed cue effects (see Supplementary Results). In response to visual stimuli, none of the additional regions were affected by the stimulus intensity or cues, except for V3A, where activity was lower following negatively skewed compared to symmetric cues (right *p* = .021, left *p* = .010).

Overall (see Supplementary Figure 1 and Supplementary Figure 2 for all effects across all regions), strong effects of stimulus intensity in pain and moderate effects in vision validated the sensitivity of ROIs to stimulation, but cue effects on stimulus-evoked activity were limited. In pain, we found aversive PE-like responses in the PAG and, in NAc, assimilation to cues and effects of cue uncertainty. However, the latter effects were not in line with previous theoretical predictions and findings ^25,70–72^. In vision, we found limited assimilation towards extreme low-value cues in some visual areas. Finally, several nociceptive regions, as well as the NPS, assimilated to the cue mean during the anticipation period, but were not affected by the cue’s precision (see Supplementary Results).

## Discussion

Current models of perception emphasize the importance of predictive processes in constructing perceptual experience across modalities. Perception has been shown to assimilate towards expected values in different modalities, such as pain ^7,8,10–12^, vision ^5,13,14^, audition ^73–75^, taste and olfaction ^76^, and interoceptive experiences like itch ^77^ and nausea ^78^. Consistent with these findings, in the current study both reported expectations and perceptions showed strong assimilation towards predicted values (cue mean) in both pain and visual intensity judgments. Furthermore, this assimilation persisted throughout the experiment although the cues were not reinforced, and was highly correlated across modalities, suggesting domain-general susceptibility to cues and/or learning. While several brain regions responded more to more intense stimuli, only neural activity related to higher-level pain processing was modulated by the cue mean. Behaviorally, in line with Bayesian predictive coding accounts ^5,6,22^, we found that overall the effect of the cue mean on perception was stronger when cue precision was higher. However, ratings were also overall higher following less precise cues, and no brain regions showed activation patterns consistent with these effects (but see below). Finally, we found that when generating expectations, people overweight extreme values across modalities, and also smaller values specifically in pain.

Our findings are mostly consistent with Bayesian predictive processing accounts, but provide new evidence addressing several open questions and potentially accounting for several previous contradicting findings. First, while the prediction regarding assimilation to expected values is straightforward at the behavioral level, where and how the brain encodes this information is much less clear. One hypothesis is that brain representations related to perceptual experience should assimilate towards predicted values. Here, early nociceptive and visual regions – and a neuromarker for nociceptive pain, the NPS – did not show evidence for assimilation towards predicted values during perception, nor did higher-level perceptual regions such as the aMCC and aIns in pain or V3, V4 and V5 in visual perception. Only the right NAc and SIIPS, a distributed neuromarker for pain related to endogenous sources (including signal in dmPFC, aIns, vmPFC, and other regions), showed significant assimilation to predicted values. These results suggest that early perceptual processes are relatively shielded from influences of conceptually driven predictions, and the strong effects on perception are driven by higher-level evaluative processes, in line with psychological accounts questioning early influences of context information on perception ^51^ and recent findings in the context of placebo analgesia ^55^. Such effects might also depend on the type of cues used. Indeed, a previous study using cues similar to the ones we have used here, has found that frontoparietal areas involved in high-level construction of perception and value (e.g., dlPFC, OFC and IPS) mediated cue effects on pain, whereas classically conditioned cue effects were mostly mediated by different systems ^10^.

Predictive processing theories also emphasize coding of PEs (‘predictive coding’ ^6,79,80^), a contrast with predictions that provide efficient representations of salient changes in the environment and drive the updating of internal models (learning) ^24,71,81^. Which areas show brain responses that assimilate to predictions or contrast with them (encode PE) is currently an active area of investigation. A working hypothesis is that early sensory areas encode sensory PEs, as they are relatively distal from internal models and shielded from their effects, whereas higher-level perceptual areas encode posterior perceptions, which represent stimuli but assimilate to predicted values. This may be particularly true when internal predictive models are conceptual, including the kinds of non-reinforced cues about others’ experience that we studied here. Here, PAG activity was higher following low vs. high cue mean, consistent with pain-related aversive PE that was found in the PAG in previous studies ^68,69^. Together with the assimilation towards predicted values in higher-level systems, our results largely support this working hypothesis, although we do not directly test encoding of different types of PEs ^80^.

Bayesian accounts of brain function also make specific predictions about the precision of expectations. More certain (precise) cues should have stronger effects on perception ^5,6,22^. Some previous studies have provided support for this hypothesis by using the precision of stimulus history as a proxy for predictive precision ^25,27^. The kinds of cues with distributions we used here offer particular advantages in studying cue precision, as the cue mean, variance (i.e., inverse precision) and other properties (e.g., extreme values) can be independently experimentally manipulated across cues. Studies using this type of cues have found results that contradict predictive processing accounts by finding null effects of cue precision ^30^ or even direct effects of predictive uncertainty on increased pain perception ^7^. In the current study, cue precision effects on perceptual ratings supported Bayesian predictive processing accounts (by showing a cue mean x variance interaction; although the effect was not significant when tested only in pain). In addition, more uncertain cues led to higher ratings, as expected by the uncertainty aversiveness account, but this effect was again significant in visual perception but not pain, and thus does not support the interpretation of Yoshida et al. ^7^, who suggested that uncertainty is aversive in the context of pain.

Several studies have recently explored the neural correlates of expectations’ precision. Predictive processing accounts predict that perceptual activity will be higher following less precise cues, because of stronger reliance on incoming sensory information. Indeed, studies have shown decreased PAG fMRI activity ^25^ and early EEG responses ^70^ for more precise expectations. Here, perceptual responses were not affected by the cue precision in both modalities, including in the PAG. The only region that was affected by the cue precision was the NAc, where activity was higher following more (rather than less) precise cues during pain perception. While this effect was in opposite direction from the effects previously shown in other (earlier) regions, it could be consistent with predictive processing, since activity in the NAc might represent the priors (expectations) and thus is expected to be stronger when the priors are more precise (and also when they are stronger, and indeed NAc activity assimilated to the cue mean).

Our design also allowed us to study effects of extreme cue values, and test how participants weight predictions of upcoming pain and visual contrast across the distribution from low to high values, which has been studied in the context of magnitude judgments ^42,43^ but not, to our knowledge, in pain. If participants over-weight extreme high-pain values – e.g., attend most to the most threatening potential values – that might explain why Yoshida et al. found increased pain with high-variance cues ^7^ while other studies have not found a main effect of uncertainty on pain ^30,35^. Here, we found that perceptual ratings were drawn towards larger extreme values, but this effect, like the effects of the cue variance, was significant mostly in visual perception and not in pain perception.

Since noxious stimuli are considered to be more ambiguous and subjective compared to visual stimuli, we expected reduced cue effects on visual perception. Indeed, computational modeling indicated that the cues were weighted more strongly in pain compared to visual perception. On the other hand, the variance and skewness of the cues mostly affected visual and not pain perception. These findings suggest that overall expectations affect pain perception more, but visual perception is more sensitive to the precision of the expectations. Finally, the persistent effects of the non-reinforced cues in both pain and visual perception might represent modulation of early perceptual processes, modulation of perceptual decision-making (higher-level affective and cognitive processes), or a post-experience report bias (i.e., demand characteristics ^82^, or conformity to alleged ratings of other participants ^83^). Our neural results suggest that cues affect perceptual decision-making, since activity in regions related to early perceptual processing was not affected, while neural activity related to higher-level processing was affected by the cues, at least in pain. Moreover, neural effects were different between the two modalities, indicating that expectation effects are modality-specific rather than modality-general, even when they operate on higher-level processes.

Beyond the effect of the cues on perception, we also studied how expectations are generated from multiple values, and anticipatory neural responses. Reported expectations assimilated towards mean values. Anticipatory responses in several pain regions, including the insula, aMCC, PAG, and even the NPS, were higher for high vs. low cue mean, while the opposite direction was found on visual trials, with lower anticipatory activity for high vs. low cue mean in several visual regions (Supplementary Results). Beyond the assimilation to the mean value, reported expectations were also biased towards extreme values in both modalities, and specifically smaller values in pain. Such effects might depend on the characteristics of the sample. The current sample consisted of healthy young adults, who may have been looking for safety signals as part of an optimism bias or may have viewed themselves as less sensitive to pain compared to others (and thus trusted lower ratings more). Putatively, different populations, such as chronic pain patients, might instead look for risk signals or view themselves as more sensitive to pain, and thus overweight larger values. Perhaps surprisingly, the tendency to overweight smaller values when forming expectations did not correlate with personality or state measures such as fear of pain, pain catastrophizing, or state anxiety. However, a growing literature suggests that task-based and self-report measures are often unrelated ^84,85^ and assess different constructs over different contexts and time scales ^86,87^.

Several limitations should be considered in the context of the current findings. First, the effects of the cue variance and skewness that were found were almost all relatively weak, and the main driver of cue effects on perceptual processing was the cue mean. Second, the inclusion of both modalities in the same experiment with interleaved blocks may have yielded cross-modality dependency that could have driven some of the results, such as the correlation between the effect of the cue mean on pain and visual ratings. Third, we did not directly compare the effects between pain and visual perception, since the scales are not directly comparable between modalities (e.g., a 10-point or a 20% increase in visual contrast rating is not necessarily perceptually equivalent to the same increase in pain rating).

Taken together, our findings suggest that perception is more complex than the recent Bayesian-driven focus on the mean and uncertainty of contextual information. More specifically, they show that extreme values have an important role in how people integrate information into expectations that later affect perception. Furthermore, these findings suggest that some aspects of expectation formation and their effect on subsequent perception are modality-general (e.g., the assimilation towards the cue mean and the importance of extreme values), while others are modality-dependent (e.g., the overweighting of smaller vs. larger values and the weight given to contextual information). Better understanding of these processes and differences between sub-populations and modalities would advance our knowledge of how people form expectations, how these expectations affect perception, and how such processes could be leveraged to improve well-being and clinical care.

## Methods

### Participants

Forty-five healthy participants (25 females; age range 18-42, mean 24.1 years) completed the study. Five additional participants were excluded because they did not complete the experiment or reported impaired vision or recent substance abuse. Participants reported no history of neurological, psychiatric, dermatologic conditions, and had not taken any medication during the 48 hours period prior to their participation. Participants reporting acute and chronic pain conditions in an initial online screening were excluded from participation. We recruited participants from the University of Colorado Boulder and local community through flyers and online ads. The Institutional Review Board of the University of Colorado Boulder approved the protocol and participants provided informed consent before the beginning of the experiment.

### Procedure Overview

Participants completed three different tasks and a set of questionnaires. First, the stimulus-response task served to estimate the stimulus-response functions for heat pain stimuli of different temperatures and flickering checkerboards of different visual contrasts. Second, in the expectation task, participants rated their expectations about heat pain and visual contrast intensities based on the cues provided in each trial without any actual stimulation. Third, in the cued-perception task, participants rated the perceived painfulness of noxious stimuli and contrask of visual stimuli that were preceded by the presentation of cues.

In session 1, the stimulus-response and expectation tasks were completed outside the MRI scanner in a behavioral laboratory. In session 2 (on average 4.4 days after session 1, range 1 - 11 days), participants first completed a battery of questionnaires on a computer, followed by a brief practice of the cued-perception task outside the scanner. Participants then moved to the MR scanner where they completed another brief practice inside the scanner to get accustomed to the trackball and the MR environment. Following a structural MRI scan, they completed the cued-perception task while their brain activity was measured with fMRI. After the cued-perception task, a functional localizer with different visual checkerboard contrasts was presented to the participants. Finally, participants completed a debriefing questionnaire outside the scanner.

### Stimulus-response task

In the first task, topical heat stimuli of five different temperatures and flickering checkerboards of five different luminance contrasts were presented to participants. After each trial, participants rated the perceived painfulness of the heat or luminance contrast of the checkerboard. The task was split into two blocks, one pain block and one vision block. In each block, 25 stimuli were presented in randomized order for a total of 50 stimuli.

Temperatures in the heat block ranged from 45-49°C, in steps of 1°C. Each temperature was used five times and each stimulation lasted for two seconds (including ramp up and down from a 32°C baseline). After a variable delay of 5 – 6 seconds, participants rated the perceived heat intensity on a computerized visual analogue scale (VAS) by moving a vertical cursor bar on the VAS. The VAS was anchored at “no pain at all” and “worst pain imaginable”. Participants were instructed to rate any heat stimulus that was noticed but not painful at the low extreme and that “worst pain imaginable” referred to the context of this experiment. They were further instructed to rate the maximum only in the case that they would have lifted the thermode to stop the experiment, which did not happen. After an inter-trial-interval (ITI) of 2 – 5 seconds, the next trial started. Before the task began, participants experienced two heat stimuli of 47°C without rating, to get familiar with the heat stimulation and reduce potential anxiety. Ratings were converted to a range between 0 - 100 in all tasks and conditions.

Black-and-white radial checkerboards were presented in block 2 at 5%, 27.5%, 50%, 72.5%, and 95% luminance contrast. Checkerboards were presented for two seconds in each trial and the contrast of each checkerboard reversed at a frequency of 8 Hz to increase responses in visual brain regions. Checkerboards covered 12° visual angle in both the behavioral and the fMRI experiment and both screens were calibrated to display the same luminance level and contrast. After each stimulus, participants rated the perceived contrast on a VAS anchored “no contrast at all” and “strongest contrast”. Similar to the pain block, an ITI of 2 – 5 sec separated two consecutive trials.

### Expectation task

In the expectation task, participants saw different distributions of intensity ratings on a computer screen and reported their expected intensity of either heat pain or luminance contrast based on these ratings. Participants were informed that the rating distributions represented the ratings of other people for heat and checkerboard stimuli similar to those they just experienced in the stimulus-response task. The ratings were marked as vertical bars on visual analogues scales as used in the stimulus-response task (see Figure 1). A set of 10 ratings was presented on a single VAS in each trial for 1.5 seconds. After a brief delay of 1.75 seconds, participants rated their expected pain intensity or contrast intensity on the same VAS as in the stimulus-response task.

The distribution of ratings shown to the participants were determined by factorial combinations of their different distribution means (five levels, *M ∈* [30, 40, 50, 60, 70]), standard deviations (two levels, *SD ∈* [5, 12.5]), and skewness (three levels: negative, positive, or symmetric). Symmetric distributions were drawn from a normal distribution and skewed distributions were drawn from a log-normal distribution. Skewness was < -0.3 for negatively skewed, > 0.3 for positively skewed, and between -0.035 to 0.035 for symmetric distributions. Here, only 9 instead of 10 elements were sampled from the log-normal distribution. A 10th element was added between 2.0 – 2.5 standard deviations above or below the mean, respectively, to ensure that each skewed distribution included at least one extreme rating. All rating distributions were re-sampled if necessary to ensure that the properties of the distribution matched the specified mean, standard deviation, and skewness, as well as the independence of the factors across trials.

Participants completed a total of 360 trials in this task split into six blocks. Participants rated heat pain expectations in three consecutive blocks and contrast intensity in the other three consecutive blocks, with the order counterbalanced across participants (three pain and then three visual or three visual and then three pain blocks). The combination of the four factors (modality, mean, standard deviation, and skewness) resulted in a total of 60 combinations. Each combination was repeated six times, once per block. The order of trials was randomized for each participant within each block.

### Cued-perception task

The cued-perception task combined the two previous tasks into a single task performed inside the MRI scanner. In each trial, participants were cued with rating distributions (as in the expectation task) for 1.5 seconds before being presented with either cutaneous heat or a flickering checkerboard for two seconds (as in the stimulus-response task). Cues and stimulation periods were separated by a brief interval of 2.5 - 5.5 seconds. Participants rated the perceived intensity of the heat or luminance contrast after another brief interval of 3 - 6 seconds. Participants had 4.5 seconds to rate the intensity using an MR compatible trackball. Trials were separated by an ITI of 3 – 6.5 seconds.

In this task, stimulation modality (heat vs. visual) and two levels of stimulation intensity were combined with different rating cue distributions similar to those used in the expectation task. The rating cues were drawn from distributions with two means (*M ∈* [30, 70]), two standard deviations (*SD ∈* [5, 12.5]), and three levels of skewness (negative skew, symmetric, positive skew). Ten ratings were presented in each trial on a VAS as described for the expectation task. The factorial combination of stimulation modality, stimulation intensity, cue distribution mean, cue distribution standard deviation, and cue distribution skew resulted in a total of 48 combinations. Each combination was presented three times to each participant resulting in a total of 144 trials split into six blocks. Each block constituted a separate fMRI recording run and included 24 trials split into two mini-blocks of 12 trials each, in which only one modality was presented. The order of modality mini-blocks and the order of trials within modality was randomized for each participant. Skin conductance was recorded during the cued-perception task. Following the cued-perception task, participants completed a visual functional localizer task, in which they were presented with a visual checkerboard with varying contrasts. Data from this task were not used in any of the analyses reported in the current paper.

### Apparatus and recordings

#### Heat stimulation

An fMRI compatible Peltier thermode (1.5 × 1.5 cm surface, PATHWAY ATS; Medoc, Inc, Israel) delivered heat to the left volar forearm of the participant in the stimulus-response task and the cued-perception task. The total stimulus duration was two seconds including ramp up and down from a 32°C baseline with extremely fast ramps (70°C/second up and 40°C/second down).

#### Visual stimulation

Radial checkerboards with a diameter of 12° visual angle flickered at a rate of 8 Hz on a 50% gray background. Screen luminance was calibrated using a Minolta LS-100 luminance meter. Stimulus presentation, thermode control, and response logging were implemented using the Psychophysics Toolbox (PTB-3, http://psychtoolbox.org/) ^88^.

### Behavioral data Analysis

Analyses were performed with R version 4.3.1 (R studio version 2023.09.0+463). For reproducibility, we used the *checkpoint* package, which installs the R packages included in the code as they were on a specific date. We set the date to April 1, 2024. 211 out of 6480 trials in the cued-perception task were excluded due to technical issues with the pain device, rating device, or MRI scanner. Additional 234 trials were excluded because the ratings were too fast to represent deliberate ratings (response time < 0.2 second) or were not completed during the 4.5 seconds long rating period (response time > 4.5 seconds).

Linear mixed-effects models were fit to the perception (stimulus-response task and cued-perception task) and the expectation (expectation task) ratings with R’s lmer function, with the packages “lme4” ^89^ and “lmerTest” ^90^. All statistical tests were two-sided. In the stimulus-response task, regressors coding modality (pain vs. vision), stimulus intensity level, and the interaction between these two variables were included in the model as fixed effects. We also fitted two additional models, one for each modality (i.e., one with pain trials only and one with visual trials only), with the stimulus intensity level as a regressor. In the expectation task, included regressors were modality, cue mean, cue variance level, and cue skewness level, with all their interactions. Again, we also fitted an additional model for each modality separately, with the same regressors (except for modality and its interactions). In the cued-perception task, we included the modality, stimulus intensity level, cue mean level, cue variance level, and cue skewness level as regressors (along with their interactions). In all tasks, when significant interaction effects were found, we tested the related simple effects. All numeric variables (ratings, intensity level for the stimulus-response task, and cue mean for the stimulus-response and expectation task) were z-scored. Binary variables (modality, cue intensity level, cue variance level, and cue mean level in the cued-perception task) were modeled with one regressor, coded as 1 (pain, or high intensity level / cue mean / cue variance level) and -1 (vision, or low intensity level / cue mean / cue variance level). The cue skewness level was modeled with two regressors using the symmetric condition as the reference.

Moreover, in all models, participants were modeled as random effects. In each model, we started with a maximal random effects structure, modeling all random effects (intercept and slopes) and their correlations ^91^. In case the model did not converge properly, we simplified the maximal model by first removing the random correlations and then reducing the random terms that indicated model converges issues (i.e., correlations of 1, or random variance of 0). Using these criteria preserves type I error while potentially increasing power when random effects estimates are near the boundary values ^92^.

### Computational modeling

#### Expectation data

We developed a computational model to test how expectations are generated from the cue data, which consists of 10 values per cue. The model assumes that in each cue, each of these 10 values is weighted based on its relative location in the distribution of the 10 values (Figure 3; the model was largely inspired by Spitzer et al., 2017 ^43^). The model yields a weight for each value, based on a combination of a power term modeling the weighting of inliers vs. outliers and a logistic term modeling the weighting of values that are smaller vs. larger than the mean. First, all 10 values of each cue were rescaled to [0,1] and then demeaned, such that cue values smaller than the mean were negative and cue values larger than the mean were positive. We denote the rescaled and demeaned value *X_i_* (where i ranges from 1 to 10 cue values).

Second, for each *X_i_*, we computed a “power term” weight with the following equation:

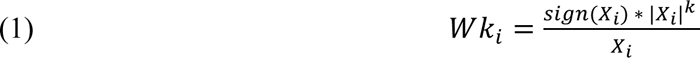

Where *Wk*_*i*_ is the power term weight of *X_i_*, and *k* ∈ (0, 1000] is a free parameter. When *k* < 1 inliers are over-weighted, when *k* = 1 outliers and inliers are equally weighted, and when *k* > 1 outliers are over-weighted. *Wk*_*i*_ values for each cue where then normalized to [0,1] by dividing each *Wk*_*i*_ by the sum of all *Wk* weights for each cue 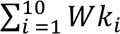.

Third, for each *X_i_*, we computed a “logistic” weight using the following equation:

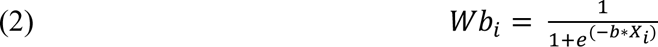

Where *Wb*_*i*_ is the logistic weight of *X_i_*, and *b* ∈ [-1000, 1000] is a free parameter. When *b* < 0 values that are smaller than the mean are over-weighted, when *b* = 0 all values are equally weighted, and when *b* > 0 values that are larger than the mean are over-weighted. *Wb*_*i*_ values are mathematically bounded between [0, 1].

Fourth, the two weights were combined, and normalized to the range [0,1], such that the weight of each *X_i_* was:

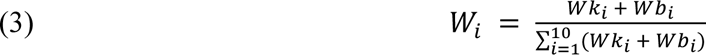

Finally, the expectation based on each cue was computed as the sum of each value *V_i_* multiplied by its weight *W_i_*:

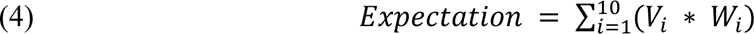

Thus, the model computes an expectation value for each cue. The model was fitted to all trials (cues) of the expectation task of each participant. The free parameters *k* and *b* were optimized per participant and modality, based on ordinary least squares (OLS), with Matlab version 2022a *lsqcurvefit* function. We then tested whether the optimized *k* was significantly different from 1 and whether the optimized *b* was significantly different from 0 at the group level, separately in each modality, with two-sided Wilcoxon signed rank tests. We also tested the correlation between the optimized *k* for pain and for vision across participants, and the same for the optimized *b*, with a two-sided Spearman correlation test.

#### Perception data

We developed five competing nested models for the cued-perception task. Models were fitted and optimized at the participant level based on OLS between the model’s predicted rating and the empirical ratings provided by the participants. Similarly to the expectation data model fitting, we used Matlab’s *lsqcurvefit* function. Then, we performed model comparison at the participant level with F tests, to test, for each participant, whether each model improves the fit compared to its nested, simpler model. To make inferences regarding the best fitting model at the group level, we performed F tests based on the mean residual sum of squares across participants, and also computed the Akaike Information Criterion (AIC) for each of the five models.

#### Model 1: Baseline Model

This model assumes participants are not affected by the cues, and their rating in each trial (*t*) depends only on the intensity of the stimulus:

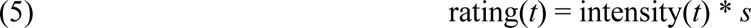

The intensity is based on the average rating of each participant for the given stimulus modality and intensity from the stimulus-response task. *s* ∈ [0, 5] is a free parameter that is used for scaling of participants’ ratings for each intensity from the stimulus-response task to the cued-perception task, which happened on a different day and in a different context (inside the MRI scanner). The Baseline Model includes two such scaling factors: *s_p_* scales the pain stimuli, and *s_v_* scales the visual stimuli. These two free parameters are included in all models.

#### Model 2: Expectation Model

This model assumes that in each trial, participants weight the cue and the stimulus to generate their rating. The weighting is determined by an additional free parameter, *w* ∈ [0, 1], that is fixed throughout the task for each participant and is the same for both modalities (thus, the model includes three free parameters):

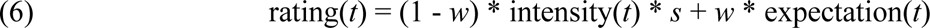

The expectation value in each trial is generated from the cue presented on that trial, with the model that was optimized for each participant based on the expectation task data (see above).

#### Model 3: Expectation Learning Model

This model adds a reinforcement learning component, such that *w* is updated based on PEs (the difference between the cue-based expectation and the rating), with learning rate *α* ∈ [-1, 1]. The rating in each trial is computed with equation 6, as in the Expectation Model, but *w* is updated in each trial based on the previous trial:

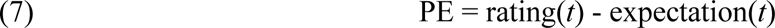

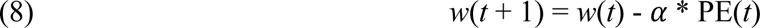

This model includes four free parameters: The two scaling factors *s_p_* and *s_v_*, the initial weighting factor *w*(*t* = 0), and the learning rate *α*.

#### Model 4: Expectation Model by Modality

This model is similar to the Expectation Model, beside one difference: the weighting factor, *w*, is separated to *w_p_* for the pain modality and *w_v_* for the visual modality. Thus, it includes four free parameters.

#### Model 5: Expectation Learning Model by Modality

This model is similar to the Expectation Learning Model, with two additions: Both the initial weighting factor (*w*(*t* = 0)) and the learning rate (*α*) are optimized separately for the pain (*w_p_* and *α_p_*) and visual (*w_v_* and *α_v_*) modalities. Thus, it includes six free parameters.

### Neuroimaging

#### Neuroimaging data acquisition

Data were collected on a 3 Tesla Siemens Trio MRI scanner with a 32 channels head coil at the University of Colorado Boulder Center for Innovation and Creativity. A high-resolution T1-weighted magnetization-prepared rapid gradient echo (MPRAGE) structural scan (0.8×0.8×0.8 mm voxels, TR: 2400 ms, TE 1: 2.07 ms, Flip angle: 8°, Tl: 1200 ms, FoV Read: 256 mm) was performed on each participant at the beginning of the MR session. We next acquired four images to compute B0-fieldmaps for distortion correction: two images with phase encoding in the anterior-posterior direction and two images with reversed phase encoding (2.7×2.7×2.7 mm voxels, TR: 7.22 s, TE: 73 ms, slices: 48, flip angle: 90°, FoV read: 220 mm). During the cued-perception task, a multiband (eight simultaneous slices) echo-planar imaging (EPI) sequence (2.7×2.7×2.7 mm voxels, TR: 410 ms, TE: 27.2 ms, slices: 48, multiband factor=8, flip angle: 44°, FoV read: 220 mm) was acquired. A single-band reference scan was acquired at the beginning of each run. We acquired 1250 volumes during each run of the cued-perception task and 776 volumes during the visual functional localizer task.

#### Neuroimaging Data preprocessing

Structural and functional data were preprocessed using fMRIPrep version 20.2.3 (RRID:SCR_016216 ^93,94^), which is based on Nipype 1.6.1 (RRID:SCR_002502 ^95,96^).

#### Anatomical data preprocessing

The T1-weighted (T1w) image was corrected for intensity non-uniformity (INU) with N4BiasFieldCorrection ^97^, distributed with ANTs 2.3.3 (RRID:SCR_004757 ^98^), and used as T1w-reference throughout the workflow. The T1w-reference was then skull-stripped with a *Nipype* implementation of the antsBrainExtraction.sh workflow (from ANTs), using OASIS30ANTs as target template. Brain tissue segmentation of cerebrospinal fluid (CSF), white-matter (WM) and gray-matter (GM) was performed on the brain-extracted T1w using fast (FSL 5.0.9, RRID:SCR_002823 ^99^). Volume-based spatial normalization to one standard space (MNI152NLin2009cAsym) was performed through nonlinear registration with antsRegistration (ANTs 2.3.3), using brain-extracted versions of both T1w reference and the T1w template. The following template was selected for spatial normalization: ICBM 152 Nonlinear Asymmetrical template version 2009c [^100^, RRID:SCR_008796; TemplateFlow ID: MNI152NLin2009cAsym],

#### Functional data preprocessing

For each of the seven BOLD runs per subject (across all tasks and sessions), the following preprocessing was performed. First, a reference volume and its skull-stripped version were generated from the single-band reference (SBRef). Susceptibility distortion correction (SDC) was omitted. The BOLD reference was then co-registered to the T1w reference using flirt (FSL 5.0.9 ^101^) with the boundary-based registration ^102^ cost-function. Co-registration was configured with nine degrees of freedom to account for distortions remaining in the BOLD reference. Head-motion parameters with respect to the BOLD reference (transformation matrices, and six corresponding rotation and translation parameters) are estimated before any spatiotemporal filtering using mcflirt (FSL 5.0.9 ^103^). First, a reference volume and its skull-stripped version were generated using a custom methodology of fMRIPrep. The BOLD time-series were resampled onto their original, native space by applying the transforms to correct for head-motion. These resampled BOLD time-series will be referred to as preprocessed BOLD in original space, or just preprocessed BOLD. The BOLD time-series were resampled into standard space, generating a preprocessed BOLD run in MNI152NLin2009cAsym space. First, a reference volume and its skull-stripped version were generated using a custom methodology of fMRIPrep. Several confounding time-series were calculated based on the preprocessed BOLD: framewise displacement (FD), DVARS and three region-wise global signals. FD was computed using two formulations following Power (absolute sum of relative motions ^104^) and Jenkinson (relative root mean square displacement between affines ^103^). FD and DVARS are calculated for each functional run, both using their implementations in Nipype (following the definitions by ^104^). All resamplings can be performed with *a single interpolation step* by composing all the pertinent transformations (i.e. head-motion transform matrices, susceptibility distortion correction when available, and co-registrations to anatomical and output spaces). Gridded (volumetric) resamplings were performed using antsApplyTransforms (ANTs), configured with Lanczos interpolation to minimize the smoothing effects of other kernels (Lanczos 1964). Non-gridded (surface) resamplings were performed using mri_vol2surf (FreeSurfer).

Many internal operations of fMRIPrep use Nilearn 0.6.2 (^105^, RRID:SCR_001362), mostly within the functional processing workflow. For more details of the pipeline, see the section corresponding to workflows in fMRIPrep’s documentation.

#### Neuroimaging data analysis

fMRI participant-level data processing was carried out using FEAT (FMRI Expert Analysis Tool) v. 6.00, part of FSL (FMRIB’s Software Library, www.fmrib.ox.ac.uk/fsl) v. 6.0.4. Data were smoothed with a Gaussian kernel of 5 mm. A 100 hz high pass filter was used during first level analysis. The first level (run level) GLM model included two regressors per trial (48 regressors of interest per run in total): separate regressors for each cue period and for each stimulation period (single trial model, or a “Beta series” ^106^). The global CSF signal and six motion parameters (translation and rotation each in three directions) were included as nuisance regressors. Variance Inflation Factor (VIF) was computed for each trial, and trials with VIF > 5 were excluded. This led to the exclusion of one trial (out of 6058 trials).

#### Neuromarker and ROI analysis

Group level analysis was performed with Matlab 2022a and 2023a, and CANlab tools (shared via Github at https://canlab.github.io/; also uses SPM12). A score for each neuromarker (NPS and SIIPS) was computed for each trial, once for the cue-evoked period and once for the stimulus-evoked period, based on the dot product between the trial’s univariate map (from the singlet trial first-level analysis) and the neuromarker weight map. Similarly, we computed for each trial the averaged activity across voxels of each of the individual a priori ROIs. For each brain measure (neuromarker or ROI), scores that were more than 3.5 SDs away from the mean score (of that same brain measure, period, and modality) were excluded. Overall, this resulted in exclusion of 0.4% of the scores (0.5% of the cue-evoked scores and 0.3% of the stimulus-evoked scores).

In order to allow comparison between the effects on the different brain measures, but keep the between-participants and within-participants effects unchanged, the scores of each brain measure were z-scored across all trials within the scope of the model (combination of modality [pain / visual perception] and period [stimulus-evoked / cue-evoked]) before including them as the dependent variable in the model (i.e., producing standardized beta estimates). Then, the z-scored scores were tested with mixed-effects models, as described above for the expectation / perceptual ratings (models were identical to the models described above, with the neuromarker / ROI score replacing the outcome rating; see Methods section *Behavioral data analysis*).

#### Multilevel mediation analysis with neuromarkers

We further tested, for each of the two neuromarkers, whether it mediated the effect of the cue-based expectation on the outcome rating, and whether it mediated the effect of the stimulus intensity on the outcome rating, each controlling for the other, for each modality separately. Brain mediation analysis tests whether a variable mediates the relationship between other variables, by identifying three statistical paths ^9^. Path *a* captures the effect of the initial variable, usually the experimental manipulation (e.g., the cue-based expectation), on the mediator (e.g., the neuromarker score). Path *b* captures the effect of the mediator on the outcome variable (e.g., the outcome rating). Path *ab* captures the indirect effect of the initial variable on the outcome variable, i.e., the part of the relationship between the initial variable and the outcome variable that is formally mediated by the mediator. Path *c’* captures the direct effect of the initial variable on the outcome variable, that is not mediated by the mediator.

Here, we tested the effect of the cue-based expectations, or the stimulus intensity level, on the outcome rating, with the neuromarker (NPS / SIIPS) as the mediator. Thus, four mediation models were tested for each modality: (1) Cue-based expectation (the trial-level expectation based on the presented cue and the expectation computational model with each participant’s optimized *k* and *b* parameters) → trial-level NPS score → trial-level outcome rating, with trial-level stimulus intensity as a covariate; (2) Trial-level stimulus intensity → trial-level NPS score → trial-level outcome rating, with trial-level cue-based expectation as a covariate; (3) Same as model 1, but with SIIPS instead of NPS; (4) Same as model 2, but with SIIPS instead of NPS. In all four models, the neuromarker score, the cue-based expectation value, and the outcome rating were z-scored across all trials, separately for each modality. Each model was run with CANlab neuroimaging analysis tools’ mediation.m function. Mediation was tested with bootstrapping, with 10000 samples ^107^.

## Data and code availability

Processed data and analysis codes are publicly shared on Github: https://github.com/rotemb9/PPRI-paper (release v1.0.0). CANlab neuroimaging analysis tools that were used as part of the analysis are available at https://canlab.github.io/. The raw imaging data will be openly shared upon publication.

## Acknowledgements

This study was funded by the National Institute of Mental Health (NIMH; R01MH076136). The funders had no role in study design, data collection and analysis, decision to publish or preparation of the manuscript. Rotem Botvinik-Nezer thanks the Golda Meir Fellowship.

## Supplementary Information

### Supplementary Results

#### Computational Modeling of Perception

We developed five computational models to test whether and how cue-based expectations are weighted when performing perception, whether the effect of expectations persist, and whether these processes are the same or different across modalities. In all models, expectations were computed for each trial of each participant based on the cue, with the optimized participant-level model from the expectation data. The intensity-based (expectation-independent) rating of each stimulus was determined based on the mean rating of each participant for a given intensity in the stimulus-response task, with a free parameter *s* per modality (*s_p_* for pain and *s_v_* for vision) that serves as a linear scaling factor, to account for the different context between the stimulus-response and cued-perception tasks (e.g., outside vs. inside the MRI scanner) and physiological processes (e.g., habituation or differences in sensitivity across skin sites).

The Baseline Model assumes the cue-based expectations do not affect perception throughout the task, and thus the rating is only based on the stimulus intensity and the scaling factor. In the Expectation Model, there are significant cue effects, and perception is based on a combination of the stimulus intensity and the cue-based expectations, with a free parameter *w* which is fixed throughout the task for each participant. This parameter determines the weight given to cue-based expectations vs. incoming information (stimulus intensity). In the “Expectation Learning Model”, there are also significant cue effects, and the relative contribution of expectation vs. stimulus intensity at the beginning of the task is determined by the free parameter *w0*. Then, *w* is updated from trial to trial based on prediction errors between the expected and empirical rating, with learning rate **α**. The two last models are equivalent to the second and third models, but include separate free parameters for each modality. In the “Expectation Model by Modality”, *w_p_* and *w_v_* model the weighting of expectations for pain and visual perception trials, respectively. In the “Expectation Learning Model by Modality”, *w0_p_* and *w0_v_* model the initial weighting of the expectation in each modality, and **α*_p_* and **α*_v_* model the learning rate for the pain and visual perception, respectively.

We fit each model to each participant’s data from the cued-perception task, and optimized the free parameters based on ordinal least squares between the predicted and empirical ratings of the painfulness or visual contrast of the stimulus. We also compared the five models and selected the best model for each participant based on F tests for nested models. Of 45 participants, the Baseline Model was the best model for 8 participants (18% of sample), the Expectation Model was best for 16 participants (35%), the Expectation Learning Model was best for 5 participants (11%), the Expectation Model by Modality was best for 8 participants (18%), and the Expectation Learning Model by Modality was best for 8 participants (18%).

In line with the participant-level results, group-level F tests for nested models based on the residual sum of squares indicated that the Expectation Model fits the data significantly better than the Baseline model (*F* = 12.40, *p* = .001), but the more complex models do not significantly improve the fit (Expectation Model vs. Expectation Learning Model: *F* = 2.04, *p* = .161; Expectation Model vs. Expectation Model by Modality: *F* = 1.06, *p* = .309; Expectation Model by Modality vs. Expectation Learning Model by Modality: *F* = 1.47, *p* = .242). Comparison based on the Akaike Information Criterion (AlC) indicated that the Expectation Model and the Expectation Learning Model were the best models while accounting for complexity (AIC values: Baseline Model = 309.81; Expectation Model = 300.17; Expectation Learning Model = 299.99 Expectation Model by Modality = 301.02; Expectation Learning Model by Modality = 301.75). These results suggest that the cues affected participants’ perception, and that only some participants learned to downweight (or ignore) the cues during the task. They further suggest that these processes were largely similar across modalities.

Finally, a direct comparison of *w_p_* and *w_v_* showed that participants weighted the cues more in pain than visual trials (paired t-test, mean difference = 0.090, *SD* difference = 0.191, *Cohen’s d* = 0.471, *t*_(44)_ = 3.162, *p* = .003), indicating greater influences of cues on pain perception than visual perception, though cue effects were strong in both modalities.

#### Effects on neural perceptual processing

In addition to the results reported in the main text, we further tested “off target” effects–stimulus intensity and cue effects on visual regions during pain perception, or pain regions during visual perception. Activity was higher (less negative) during more intense heat stimuli in several visual regions, probably because noxious stimuli produce diffuse effects, but none of them was affected by the cue during pain perception. Activity in the right IPS was higher during pain perception following cues with lower mean (similar to the aversive PE-like activity in the PAG; *p* = .047). Conversely, activity during visual perception was not affected by the stimulus intensity in any of the pain processing regions (all *p*s ≥ .162), but it was higher following cues with lower mean in the right VPL/VPM thalamus (*p* = .046), and lower following negatively skewed compared to symmetric cues in the right dorsal posterior insula (*p* = .001) and right S1 (*p* = .018). In addition, there was a significant cue mean x cue variance interaction effect on activity in the right NAc (shell-like) during visual perception (*p* = .048), such that activity was slightly more negative following high compared to low mean for low variance cues, but less negative for high compared to low mean for high variance cues (not in line with Bayesian predictive coding accounts).

#### Effects on anticipatory neural activity

While both the current study and most previous ones focused on cue effects on stimulus-evoked activity, we also performed an exploratory analysis to test anticipatory activity (see Supplementary Figure 3 and Supplementary Figure 4). Several pain processing regions were more active during anticipation in response to high vs. low cue mean, including the dorsal posterior insula (right *p* = .030; left *p* = .039), aMCC (*p* = .008), PAG (*p* = .049), and anterior insula (right *p* = .023; left *p* = .010). In addition, cue-evoked activity in the PAG (*p* = .043) and NAc (core-like; right *p* = .020; left *p* = .049) was lower following positively skewed compared to symmetric cues (opposite direction than expected and observed behaviorally, although it was not significant behaviorally). On visual trials, anticipatory activity was lower following high vs. low cue mean in V4 (right *p* = .025; left *p* = .028), left V5 (*p* = .001), left VMV2 (*p* = .003), left VMV3 (*p* = .017), left V3B (*p* = .005), left V4t (*p* = .001), IPS (right *p* = .001; left *p* = .034), and left dlPFC (*p* = .046). It was also higher for cues with higher variance (less certain cues) in right V4 (*p* = .020), left NAc (core-like, *p* = .017) and VPL/VPM thalamus (right *p* = .046, left *p* = .038). As for the skewness, anticipatory activity in visual trials was higher for negatively skewed compared to symmetric cues (opposite from the behavioral direction) in the right IPS (*p* = .006) and lower for positively skewed compared to symmetric cues (again, opposite from the behavioral effect) in the right NAc (core-like *p* = .004; shell-like *p* = .018). Finally, there was a significant interaction between the cue mean and the cue variance in the right VMV3 (*p* = .042), such that anticipatory activity was higher for high compared to low cue mean for more certain cues, and higher for low compared to high cue mean for less certain cues (not in line with Bayesian predictive coding accounts).

**Supplementary Figure 1.**
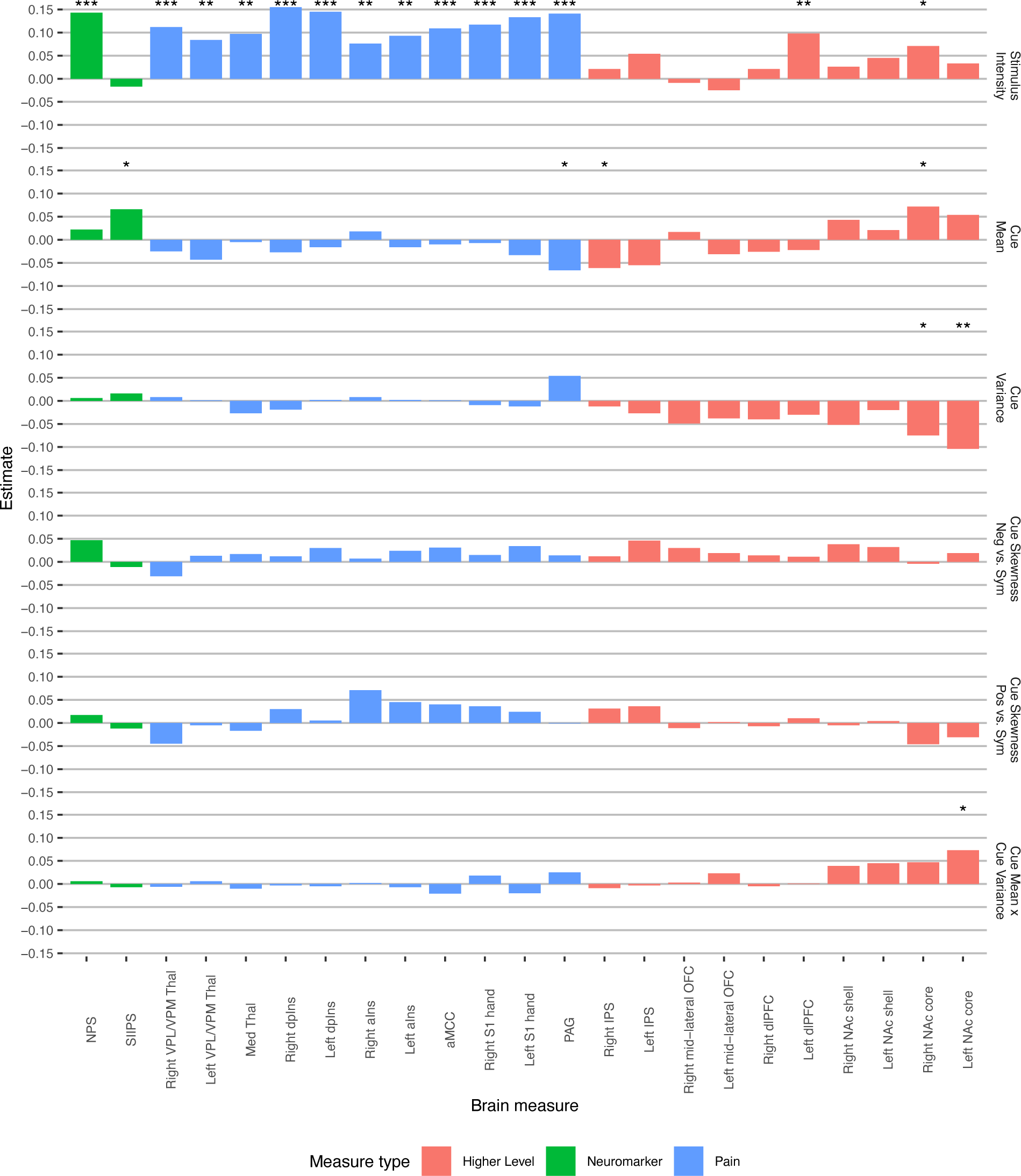
Stimulus intensity and cue effects across all ROIs during thermal stimuli. Asterisks represent the level of significance (* *p* < .05, ** *p* < .01, *** *p* < .001). Abbreviations: Neg = negative; Pos = positive; Sym = symmetric; NPS = Neurological Brain Signature; SIIPS = Stimulus Intensity Independent Pain Signature; VPL/VPM = ventral posterior lateral/medial; Thal = thalamus; dpIns = dorsal posterior insula; aIns = anterior insula; aMCC = anterior midcingulate cortex; PAG = periaqueductal gray; IPS = intraparietal sulcus; OFC = orbitofrontal cortex; dlPFC = dorsolateral prefrontal cortex; NAc = nucleus accumbens.

**Supplementary Figure 2.**
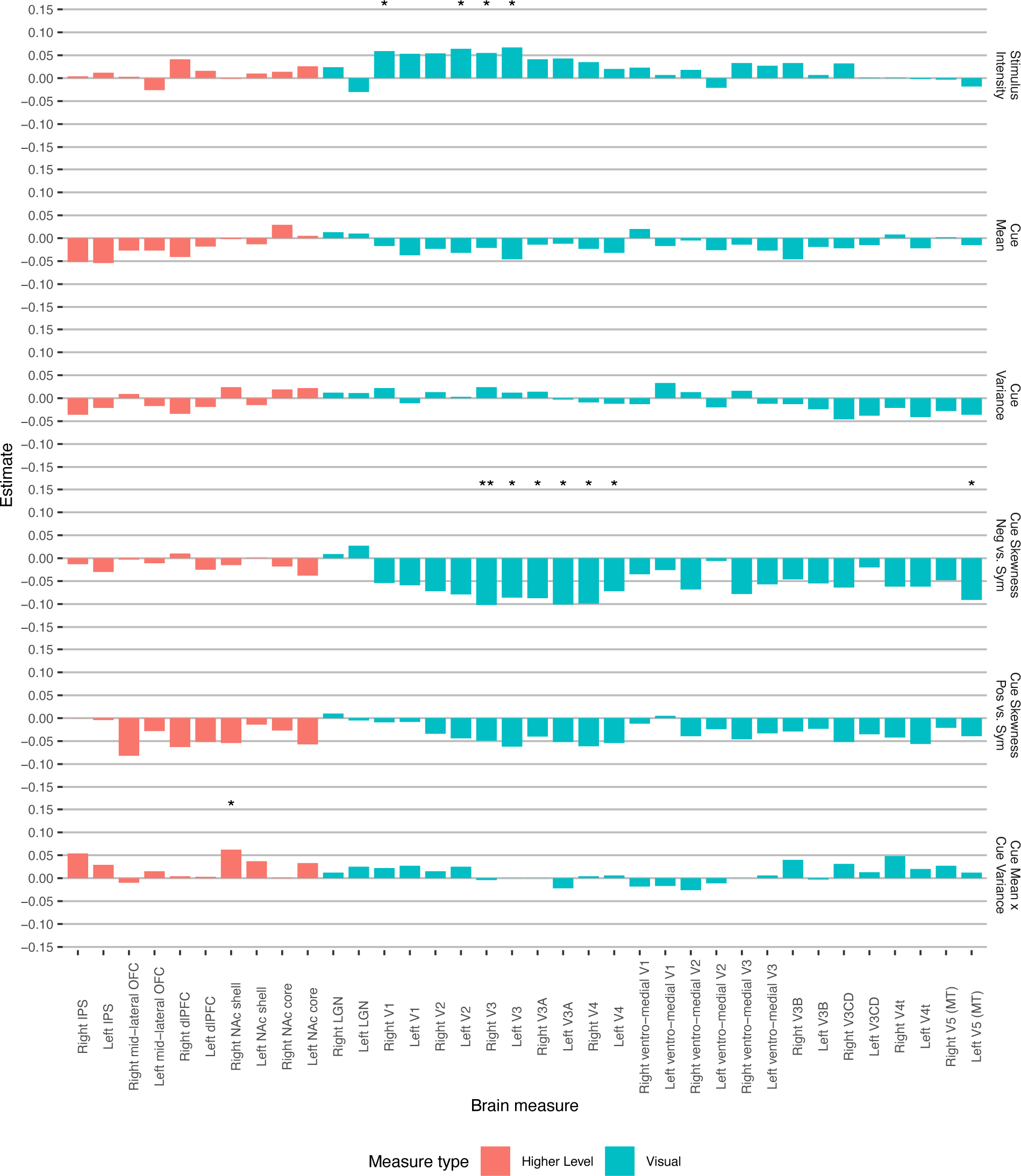
Stimulus intensity and cue effects across all ROIs during visual stimuli. Asterisks represent the level of significance (* *p* < .05, ** *p* < .01, *** *p* < .001). Abbreviations: Neg = negative; Pos = positive; Sym = symmetric; IPS = intraparietal sulcus; OFC = orbitofrontal cortex; dlPFC = dorsolateral prefrontal cortex; NAc = nucleus accumbens; LGN = lateral geniculate nucleus.

**Supplementary Figure 3.**
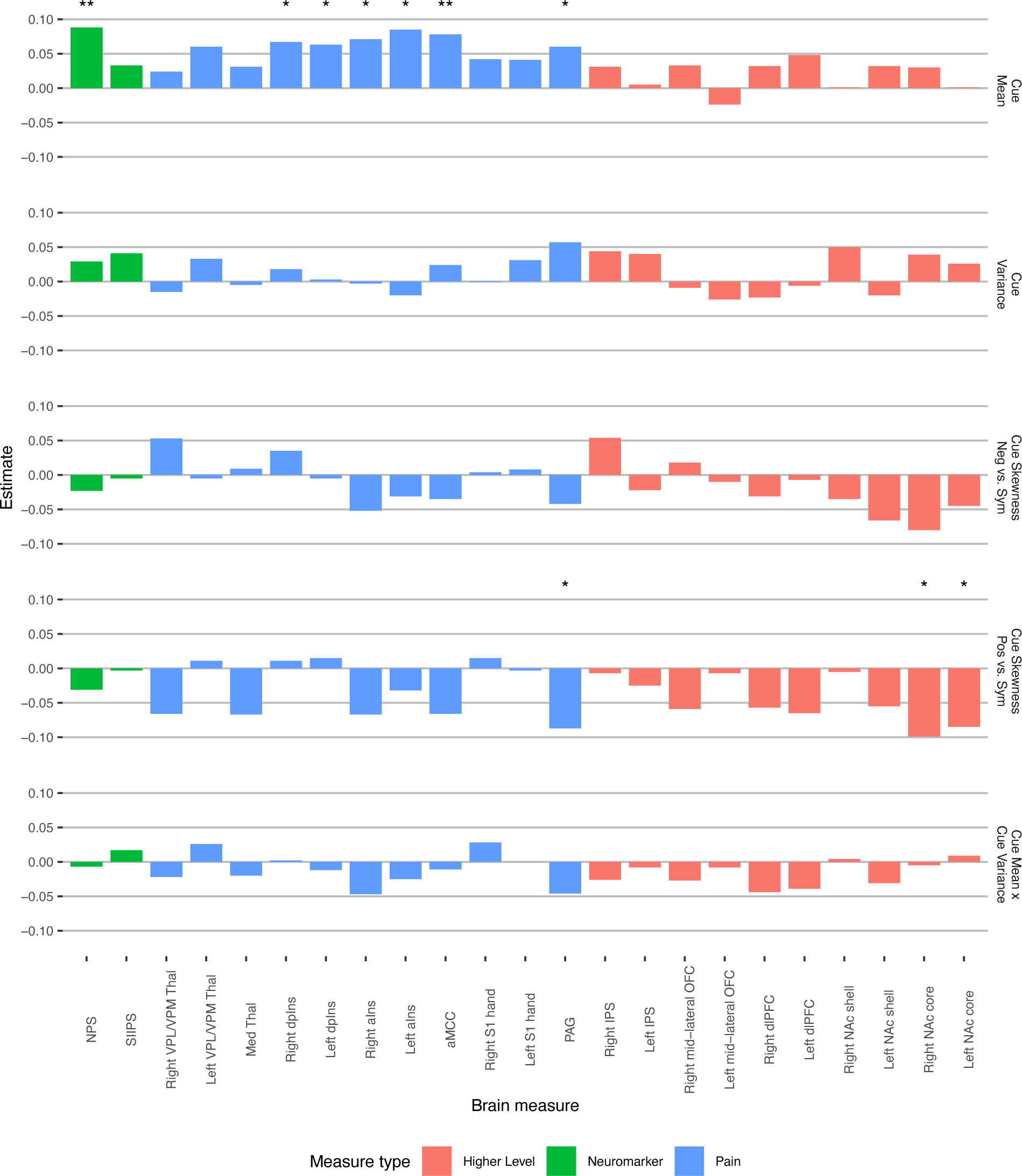
Cue effects across all ROIs during anticipation for thermal stimuli. Asterisks represent the level of significance (* *p* < .05, ** *p* < .01, *** *p* < .001). Abbreviations: Neg = negative; Pos = positive; Sym = symmetric; NPS = Neurological Brain Signature; SIIPS = Stimulus Intensity Independent Pain Signature; VPL/VPM = ventral posterior lateral/medial; Thal = thalamus; dpIns = dorsal posterior insula; aIns = anterior insula; aMCC = anterior midcingulate cortex; PAG = periaqueductal gray; IPS = intraparietal sulcus; OFC = orbitofrontal cortex; dlPFC = dorsolateral prefrontal cortex; NAc = nucleus accumbens.

**Supplementary Figure 4.**
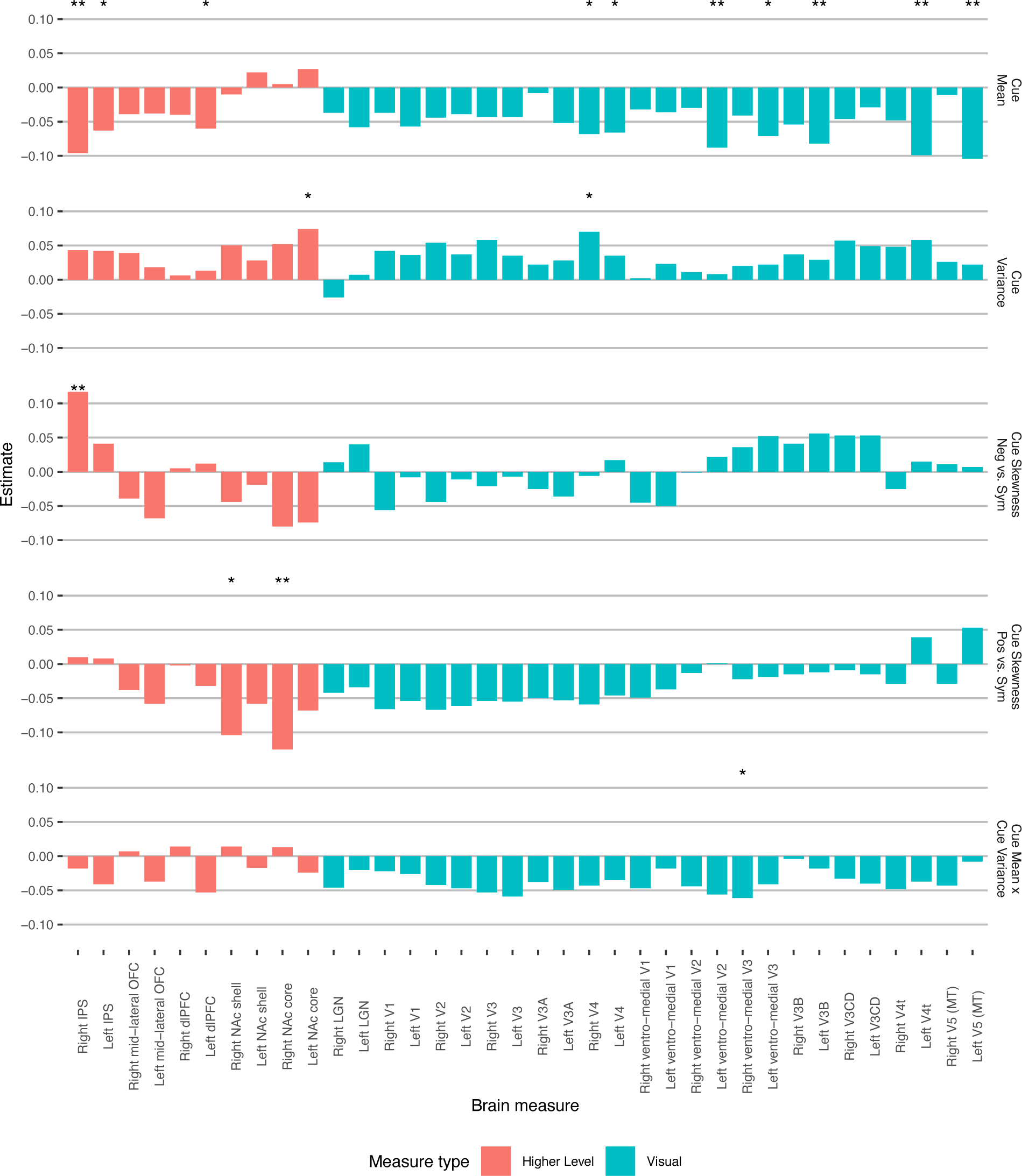
Cue effects across all ROIs during anticipation for visual stimuli. Asterisks represent the level of significance (* *p* < .05, ** *p* < .01, *** *p* < .001). Abbreviations: Neg = negative; Pos = positive; Sym = symmetric; IPS = intraparietal sulcus; OFC = orbitofrontal cortex; dlPFC = dorsolateral prefrontal cortex; NAc = nucleus accumbens; LGN = lateral geniculate nucleus.

**Supplementary Table 1.**
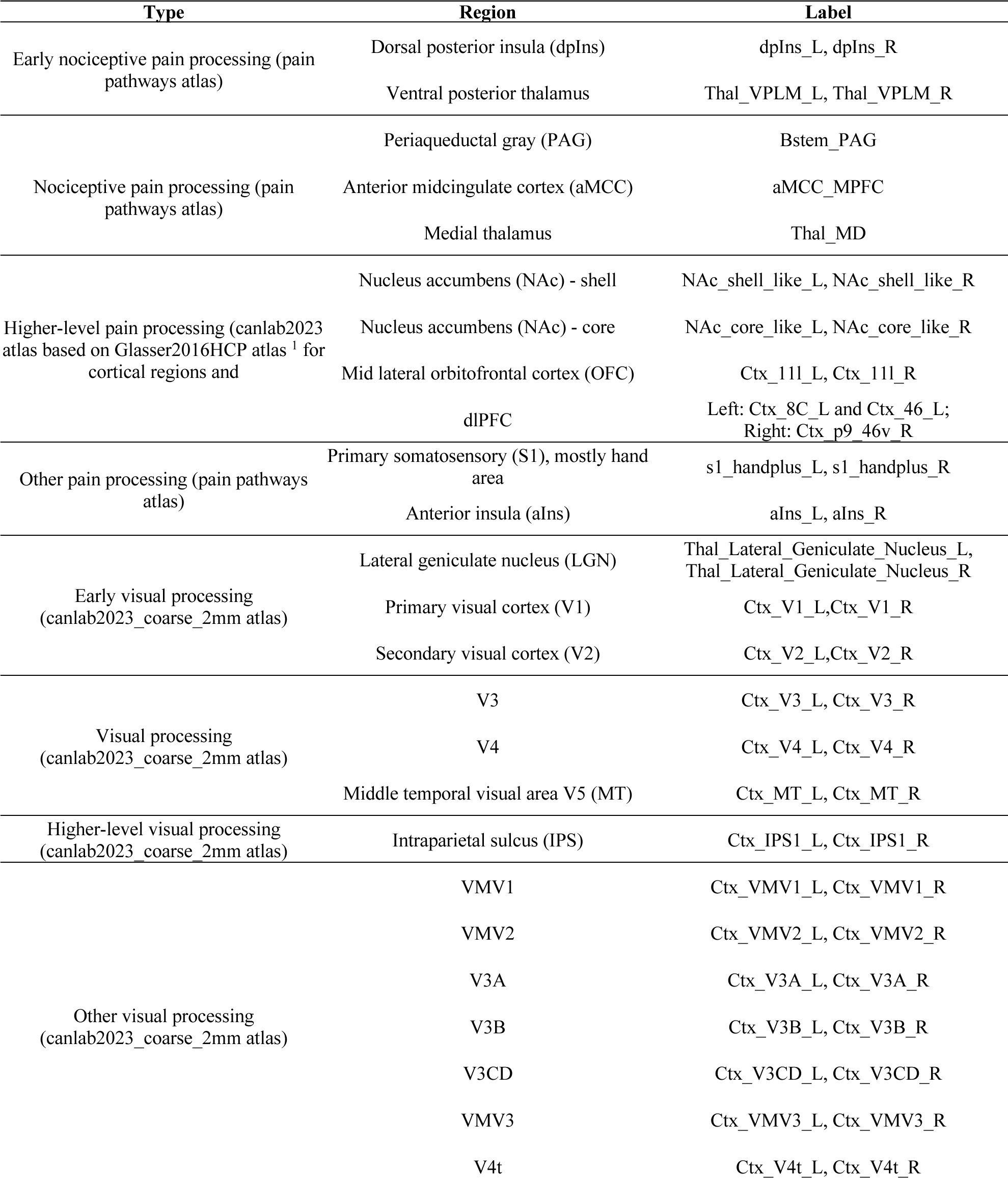
A priori regions of interest. Atlases are available via CANlab neuroimaging analysis tools (https://canlab.github.io/). The canlab2023 atlas is used with threshold = 0.2, and its regions included here are based on Glasser2016HCP atlas ^1^ for cortical regions, Tian2020 atlas ^2^ for the NAc, and Morel2010 atlas ^3^ for the thalamus.

**Supplementary Table 2.** Statistics for the effects on NPS and SIIPS scores during visual perception.

| Neuromarker | Effect | Estimate | SE | DF | t | p |
| --- | --- | --- | --- | --- | --- | --- |
| NPS | Stimulus Intensity | -0.015 | 0.030 | 2947.1 | -0.50 | 0.617 |
|  | Cue mean | -0.013 | 0.030 | 2946.4 | -0.45 | 0.652 |
|  | Cue var | 0.022 | 0.030 | 2947.9 | 0.75 | 0.456 |
|  | Cue SK (neg) | -0.055 | 0.042 | 2947.4 | -1.29 | 0.196 |
|  | Cue SK (pos) | 0.038 | 0.042 | 2947.3 | 0.90 | 0.368 |
|  | Cue mean x cue var | 0.000 | 0.030 | 2946.4 | 0.02 | 0.988 |
| SIIPS | Stimulus Intensity | -0.027 | 0.030 | 2935.7 | -0.88 | 0.377 |
|  | Cue mean | -0.005 | 0.030 | 2935.1 | -0.17 | 0.864 |
|  | Cue var | 0.051 | 0.030 | 2936.7 | 1.68 | 0.094 |
|  | Cue SK (neg) | 0.008 | 0.043 | 2936.1 | 0.20 | 0.844 |
|  | Cue SK (pos) | 0.035 | 0.043 | 2935.9 | 0.81 | 0.417 |
|  | Cue mean x cue var | -0.011 | 0.030 | 2934.9 | -0.38 | 0.704 |

**Supplementary Table 3.** Statistics for the effects on ROIs related to early pain perceptual processing. Abbreviations: var = variance; SK = skewness; neg = negative (vs. symmetric); pos = positive (vs. symmetric); stim int = stimulus intensity; VPL/VPM Thal = ventral posterior thalamus (medial and lateral); dpIns = dorsal posterior insula. Significant effects (uncorrected *p* < .05) are marked in bold.

| Region | Effect | Estimate | SE | DF | t | p |
| --- | --- | --- | --- | --- | --- | --- |
| Left<br>dpIns | **Stimulus intensity** | **0.145** | **0.028** | **2906.0** | **5.17** | **< .001** |
|  | Cue mean | -0.016 | 0.029 | 302.1 | -0.55 | 0.585 |
|  | Cue var | 0.002 | 0.028 | 2905.7 | 0.06 | 0.956 |
|  | Cue SK (neg) | 0.030 | 0.040 | 2905.3 | 0.76 | 0.444 |
|  | Cue SK (pos) | 0.005 | 0.040 | 2906.3 | 0.14 | 0.892 |
|  | Cue mean x cue var | -0.005 | 0.028 | 2913.9 | -0.18 | 0.858 |
|  | Stim int x cue mean | -0.001 | 0.028 | 2904.7 | -0.05 | 0.963 |
| Left<br>VPL/VPM<br>Thal | **Stimulus intensity** | **0.084** | **0.029** | **2906.8** | **2.93** | **0.003** |
|  | Cue mean | -0.043 | 0.029 | 314.4 | -1.47 | 0.141 |
|  | Cue var | 0.001 | 0.029 | 2906.6 | 0.05 | 0.962 |
|  | Cue SK (neg) | 0.013 | 0.040 | 2906.1 | 0.31 | 0.757 |
|  | Cue SK (pos) | -0.005 | 0.040 | 2907.2 | -0.13 | 0.898 |
|  | Cue mean x cue var | 0.006 | 0.029 | 2914.8 | 0.22 | 0.830 |
|  | Stim int x cue mean | 0.005 | 0.029 | 2905.6 | 0.17 | 0.862 |
| Right<br>dpIns | **Stimulus intensity** | **0.155** | **0.027** | **2948.3** | **5.67** | **< .001** |
|  | Cue mean | -0.027 | 0.027 | 2948.6 | -1.00 | 0.316 |
|  | Cue var | -0.019 | 0.027 | 2949.1 | -0.68 | 0.499 |
|  | Cue SK (neg) | 0.012 | 0.039 | 2948.6 | 0.31 | 0.754 |
|  | Cue SK (pos) | 0.030 | 0.039 | 2948.8 | 0.78 | 0.435 |
|  | Cue mean x cue var | -0.003 | 0.027 | 2948.5 | -0.10 | 0.916 |
|  | Stim int x cue mean | -0.007 | 0.027 | 2948.7 | -0.24 | 0.808 |
| Right<br>VPL/VPM<br>Thal | **Stimulus intensity** | **0.112** | **0.029** | **2946.5** | **3.88** | **< .001** |
|  | Cue mean | -0.025 | 0.029 | 2946.9 | -0.87 | 0.385 |
|  | Cue var | 0.008 | 0.029 | 2947.7 | 0.28 | 0.783 |
|  | Cue SK (neg) | -0.031 | 0.041 | 2946.9 | -0.75 | 0.456 |
|  | Cue SK (pos) | -0.045 | 0.041 | 2947.3 | -1.10 | 0.273 |
|  | Cue mean x cue var | -0.006 | 0.029 | 2946.8 | -0.21 | 0.836 |
|  | Stim int x cue mean | -0.027 | 0.029 | 2947.0 | -0.94 | 0.346 |

**Supplementary Table 4.** Statistics for the effects on ROIs related to early visual perceptual processing. Abbreviations: var = variance; SK = skewness; neg = negative (vs. symmetric); pos = positive (vs. symmetric); stim int = stimulus intensity; LGN = lateral geniculate nucleus. Significant effects (uncorrected *p* < .05) are marked in bold.

| Region | Effect | Estimate | SE | DF | t | p |
| --- | --- | --- | --- | --- | --- | --- |
| Right LGN | Stimulus Intensity | 0.024 | 0.031 | 2926.4 | 0.78 | 0.438 |
|  | Cue mean | 0.013 | 0.031 | 308.5 | 0.41 | 0.683 |
|  | Cue var | 0.012 | 0.031 | 2926.1 | 0.40 | 0.687 |
|  | Cue SK (neg) | 0.009 | 0.043 | 2926.0 | 0.21 | 0.837 |
|  | Cue SK (pos) | 0.010 | 0.043 | 2925.8 | 0.23 | 0.819 |
|  | Cue mean x var | 0.012 | 0.031 | 2927.4 | 0.38 | 0.702 |
|  | Stim int x cue mean | -0.014 | 0.031 | 2927.1 | -0.46 | 0.646 |
| Left LGN | Stimulus Intensity | -0.030 | 0.031 | 2963.7 | -0.97 | 0.332 |
|  | Cue mean | 0.010 | 0.031 | 2962.8 | 0.34 | 0.736 |
|  | Cue var | 0.011 | 0.031 | 2963.8 | 0.37 | 0.712 |
|  | Cue SK (neg) | 0.027 | 0.044 | 2963.3 | 0.63 | 0.531 |
|  | Cue SK (pos) | -0.005 | 0.043 | 2963.7 | -0.12 | 0.902 |
|  | Cue mean x var | 0.025 | 0.031 | 2962.7 | 0.81 | 0.418 |
|  | Stim int x cue mean | -0.023 | 0.031 | 2962.8 | -0.75 | 0.456 |
| Right V1 | **Stimulus Intensity** | **0.059** | **0.029** | **2947.0** | **2.04** | **0.041** |
|  | Cue mean | -0.017 | 0.029 | 2946.4 | -0.58 | 0.563 |
|  | Cue var | 0.022 | 0.029 | 2947.0 | 0.78 | 0.438 |
|  | Cue SK (neg) | -0.054 | 0.041 | 2946.8 | -1.32 | 0.187 |
|  | Cue SK (pos) | -0.009 | 0.041 | 2946.6 | -0.23 | 0.821 |
|  | Cue mean x var | 0.022 | 0.029 | 2946.3 | 0.75 | 0.456 |
|  | Stim int x cue mean | -0.032 | 0.029 | 2946.4 | -1.11 | 0.269 |
| Left V1 | Stimulus Intensity | 0.053 | 0.029 | 2951.0 | 1.83 | 0.067 |
|  | Cue mean | -0.037 | 0.029 | 2950.4 | -1.27 | 0.204 |
|  | Cue var | -0.011 | 0.029 | 2951.0 | -0.39 | 0.696 |
|  | Cue SK (neg) | -0.059 | 0.041 | 2950.8 | -1.43 | 0.153 |
|  | Cue SK (pos) | -0.008 | 0.041 | 2950.6 | -0.21 | 0.838 |
|  | Cue mean x var | 0.027 | 0.029 | 2950.4 | 0.94 | 0.348 |
|  | Stim int x cue mean | -0.031 | 0.029 | 2950.4 | -1.06 | 0.289 |
| Right V2 | Stimulus Intensity | 0.054 | 0.028 | 2956.6 | 1.89 | 0.058 |
|  | Cue mean | -0.023 | 0.028 | 2956.1 | -0.80 | 0.423 |
|  | Cue var | 0.013 | 0.028 | 2956.5 | 0.46 | 0.646 |
|  | Cue SK (neg) | -0.072 | 0.040 | 2956.4 | -1.80 | 0.072 |
|  | Cue SK (pos) | -0.034 | 0.040 | 2956.3 | -0.84 | 0.401 |
|  | Cue mean x var | 0.015 | 0.028 | 2956.0 | 0.51 | 0.608 |
|  | Stim int x cue mean | -0.021 | 0.028 | 2956.1 | -0.74 | 0.461 |
| Left V2 | **Stimulus Intensity** | **0.064** | **0.029** | **2953.0** | **2.20** | **0.028** |
|  | Cue mean | -0.032 | 0.029 | 2952.4 | -1.09 | 0.275 |
|  | Cue var | 0.003 | 0.029 | 2953.0 | 0.11 | 0.916 |
|  | Cue SK (neg) | -0.079 | 0.041 | 2952.8 | -1.92 | 0.056 |
|  | Cue SK (pos) | -0.044 | 0.041 | 2952.6 | -1.08 | 0.282 |
|  | Cue mean x var | 0.025 | 0.029 | 2952.4 | 0.85 | 0.395 |
|  | Stim int x cue mean | -0.021 | 0.029 | 2952.4 | -0.74 | 0.460 |

**Supplementary Table 5.** Statistics for the effects on ROIs related to pain perceptual processing. Abbreviations: var = variance; SK = skewness; neg = negative (vs. symmetric); pos = positive (vs. symmetric); stim int = stimulus intensity; aMCC = anterior midcingulate cortex; Med Thal = medial thalamus; PAG = periaqueductal gray. Significant effects (uncorrected *p* < .05) are marked in bold.

| Region | Effect | Estimate | SE | DF | t | p |
| --- | --- | --- | --- | --- | --- | --- |
| aMCC | **Stimulus Intensity** | **0.109** | **0.028** | **2905.4** | **3.92** | **< .001** |
|  | Cue mean | -0.010 | 0.029 | 243.7 | -0.35 | 0.728 |
|  | Cue var | 0.001 | 0.028 | 2905.0 | 0.03 | 0.973 |
|  | Cue SK (neg) | 0.031 | 0.039 | 2904.8 | 0.79 | 0.432 |
|  | Cue SK (pos) | 0.040 | 0.039 | 2905.5 | 1.03 | 0.303 |
|  | Cue mean x var | -0.021 | 0.028 | 2912.4 | -0.75 | 0.453 |
|  | Stim int x cue mean | -0.029 | 0.028 | 2904.1 | -1.06 | 0.291 |
| Med Thal | **Stimulus Intensity** | **0.097** | **0.029** | **2948.5** | **3.34** | **0.001** |
|  | Cue mean | -0.005 | 0.029 | 2948.9 | -0.16 | 0.872 |
|  | Cue var | -0.027 | 0.029 | 2949.8 | -0.94 | 0.347 |
|  | Cue SK (neg) | 0.017 | 0.041 | 2949.0 | 0.42 | 0.674 |
|  | Cue SK (pos) | -0.017 | 0.041 | 2949.4 | -0.41 | 0.680 |
|  | Cue mean x var | -0.010 | 0.029 | 2948.8 | -0.35 | 0.728 |
|  | Stim int x cue mean | -0.010 | 0.029 | 2949.1 | -0.35 | 0.726 |
| PAG | **Stimulus Intensity** | **0.141** | **0.030** | **2907.3** | **4.71** | **< .001** |
|  | **Cue mean** | **-0.066** | **0.030** | **341.9** | **-2.17** | **0.031** |
|  | Cue var | 0.054 | 0.030 | 2907.8 | 1.79 | 0.073 |
|  | Cue SK (neg) | 0.014 | 0.042 | 2906.8 | 0.33 | 0.739 |
|  | Cue SK (pos) | -0.001 | 0.042 | 2907.9 | -0.04 | 0.972 |
|  | Cue mean x var | 0.025 | 0.030 | 2915.3 | 0.83 | 0.406 |
|  | Stim int x cue mean | -0.028 | 0.030 | 2906.4 | -0.92 | 0.357 |

**Supplementary Table 6.** Statistics for the effects on ROIs related to visual perceptual processing. Abbreviations: var = variance; SK = skewness; neg = negative (vs. symmetric); pos = positive (vs. symmetric); stim int = stimulus intensity; MT = middle temporal. Significant effects (uncorrected *p* < .05) are marked in bold.

| Region | Effect | Estimate | SE | DF | t | p |
| --- | --- | --- | --- | --- | --- | --- |
| Right V3 | **Stimulus Intensity** | **0.055** | **0.027** | **2961.3** | **2.00** | **0.046** |
|  | Cue mean | -0.021 | 0.027 | 2960.9 | -0.77 | 0.439 |
|  | Cue var | 0.024 | 0.027 | 2961.3 | 0.87 | 0.386 |
|  | **Cue SK (neg)** | **-0.102** | **0.039** | **2961.2** | **-2.63** | **0.009** |
|  | Cue SK (pos) | -0.049 | 0.039 | 2961.1 | -1.26 | 0.207 |
|  | Cue mean x var | -0.004 | 0.027 | 2960.9 | -0.13 | 0.897 |
|  | Stim int x cue mean | -0.018 | 0.027 | 2960.9 | -0.67 | 0.505 |
| Left V3 | **Stimulus Intensity** | **0.067** | **0.028** | **2955.9** | **2.39** | **0.017** |
|  | Cue mean | -0.046 | 0.028 | 2955.5 | -1.62 | 0.106 |
|  | Cue var | 0.012 | 0.028 | 2955.9 | 0.42 | 0.672 |
|  | **Cue SK (neg)** | **-0.086** | **0.040** | **2955.7** | **-2.15** | **0.032** |
|  | Cue SK (pos) | -0.062 | 0.040 | 2955.6 | -1.54 | 0.123 |
|  | Cue mean x var | 0.000 | 0.028 | 2955.4 | -0.01 | 0.993 |
|  | Stim int x cue mean | -0.014 | 0.028 | 2955.4 | -0.48 | 0.631 |
| Right V4 | Stimulus Intensity | 0.035 | 0.027 | 2953.5 | 1.30 | 0.193 |
|  | Cue mean | -0.023 | 0.027 | 2953.2 | -0.84 | 0.402 |
|  | Cue var | -0.009 | 0.027 | 2953.6 | -0.32 | 0.747 |
|  | **Cue SK (neg)** | **-0.099** | **0.038** | **2953.5** | **-2.58** | **0.010** |
|  | Cue SK (pos) | -0.061 | 0.038 | 2953.4 | -1.59 | 0.111 |
|  | Cue mean x var | 0.004 | 0.027 | 2953.2 | 0.13 | 0.894 |
|  | Stim int x cue mean | -0.005 | 0.027 | 2953.2 | -0.18 | 0.857 |
| Left V4 | Stimulus Intensity | 0.020 | 0.025 | 2959.5 | 0.81 | 0.421 |
|  | Cue mean | -0.032 | 0.025 | 2959.2 | -1.28 | 0.201 |
|  | Cue var | -0.012 | 0.025 | 2959.5 | -0.49 | 0.624 |
|  | **Cue SK (neg)** | **-0.072** | **0.036** | **2959.4** | **-2.01** | **0.045** |
|  | Cue SK (pos) | -0.054 | 0.036 | 2959.3 | -1.52 | 0.130 |
|  | Cue mean x var | 0.006 | 0.025 | 2959.2 | 0.25 | 0.805 |
|  | Stim int x cue mean | 0.008 | 0.025 | 2959.2 | 0.33 | 0.740 |
| Right V5 (MT) | Stimulus Intensity | -0.003 | 0.029 | 2957.0 | -0.10 | 0.922 |
|  | Cue mean | 0.002 | 0.029 | 2956.5 | 0.06 | 0.954 |
|  | Cue var | -0.028 | 0.029 | 2957.0 | -0.99 | 0.321 |
|  | Cue SK (neg) | -0.048 | 0.041 | 2956.9 | -1.17 | 0.242 |
|  | Cue SK (pos) | -0.021 | 0.041 | 2956.8 | -0.51 | 0.608 |
|  | Cue mean x var | 0.027 | 0.029 | 2956.5 | 0.94 | 0.349 |
|  | Stim int x cue mean | -0.008 | 0.029 | 2956.5 | -0.27 | 0.784 |
| Left V5 (MT) | Stimulus Intensity | -0.018 | 0.029 | 2961.1 | -0.63 | 0.531 |
|  | Cue mean | -0.015 | 0.029 | 2960.7 | -0.51 | 0.607 |
|  | Cue var | -0.036 | 0.029 | 2961.2 | -1.24 | 0.215 |
|  | **Cue SK (neg)** | **-0.091** | **0.041** | **2961.0** | **-2.22** | **0.027** |
|  | Cue SK (pos) | -0.039 | 0.041 | 2960.9 | -0.95 | 0.344 |
|  | Cue mean x var | 0.012 | 0.029 | 2960.6 | 0.43 | 0.666 |
|  | Stim int x cue mean | 0.044 | 0.029 | 2960.7 | 1.53 | 0.126 |

**Supplementary Table 7.** Statistics for the effects on ROIs related to higher level pain processing. Abbreviations: var = variance; SK = skewness; neg = negative (vs. symmetric); pos = positive (vs. symmetric); stim int = stimulus intensity; dlPFC = dorsolateral prefrontal cortex; OFC = orbitofrontal cortex; NAc = nucleus accumbens. Significant effects (uncorrected *p* < .05) are marked in bold.

| Region | Effect | Estimate | SE | DF | t | p |
| --- | --- | --- | --- | --- | --- | --- |
| Right dlPFC | Stimulus Intensity | 0.021 | 0.03 | 2948.4 | 0.72 | 0.473 |
|  | Cue mean | -0.026 | 0.03 | 2948.8 | -0.87 | 0.386 |
|  | Cue var | -0.04 | 0.03 | 2949.9 | -1.34 | 0.181 |
|  | Cue SK (neg) | 0.014 | 0.042 | 2949 | 0.34 | 0.737 |
|  | Cue SK (pos) | -0.007 | 0.042 | 2949.4 | -0.17 | 0.868 |
|  | Cue mean x var | -0.005 | 0.03 | 2948.7 | -0.18 | 0.854 |
|  | Stim int x cue mean | 0.024 | 0.03 | 2949 | 0.81 | 0.417 |
| Left dlPFC | **Stimulus Intensity** | **0.098** | **0.03** | **2948.5** | **3.31** | **0.001** |
|  | Cue mean | -0.022 | 0.03 | 2948.9 | -0.76 | 0.447 |
|  | Cue var | -0.03 | 0.03 | 2950 | -1.02 | 0.307 |
|  | Cue SK (neg) | 0.011 | 0.042 | 2949 | 0.26 | 0.798 |
|  | Cue SK (pos) | 0.01 | 0.042 | 2949.5 | 0.25 | 0.801 |
|  | Cue mean x var | 0.001 | 0.03 | 2948.8 | 0.05 | 0.961 |
|  | Stim int x cue mean | 0.014 | 0.03 | 2949.1 | 0.48 | 0.631 |
| Right mid-lateral OFC | Stimulus Intensity | -0.009 | 0.03 | 2948.5 | -0.31 | 0.757 |
|  | Cue mean | 0.017 | 0.03 | 2948.9 | 0.58 | 0.565 |
|  | Cue var | -0.049 | 0.03 | 2950 | -1.64 | 0.102 |
|  | Cue SK (neg) | 0.03 | 0.042 | 2949 | 0.72 | 0.469 |
|  | Cue SK (pos) | -0.011 | 0.042 | 2949.5 | -0.26 | 0.793 |
|  | Cue mean x var | 0.003 | 0.03 | 2948.8 | 0.11 | 0.916 |
|  | Stim int x cue mean | 0.01 | 0.03 | 2949.1 | 0.32 | 0.748 |
| Left mid-lateral OFC | Stimulus Intensity | -0.025 | 0.03 | 2904.5 | -0.85 | 0.394 |
|  | Cue mean | -0.031 | 0.031 | 292.8 | -1.02 | 0.308 |
|  | Cue var | -0.038 | 0.03 | 2904.6 | -1.29 | 0.198 |
|  | Cue SK (neg) | 0.019 | 0.042 | 2903.9 | 0.46 | 0.647 |
|  | Cue SK (pos) | 0.002 | 0.042 | 2904.9 | 0.04 | 0.97 |
|  | Cue mean x var | 0.023 | 0.03 | 2911.6 | 0.79 | 0.432 |
|  | Stim int x cue mean | 0.028 | 0.03 | 2903.4 | 0.95 | 0.342 |
| Right NAc core | **Stimulus Intensity** | **0.071** | **0.029** | **2904.9** | **2.39** | **0.017** |
|  | **Cue mean** | **0.072** | **0.03** | **315.4** | **2.39** | **0.018** |
|  | **Cue var** | **-0.075** | **0.029** | **2905** | **-2.53** | **0.011** |
|  | Cue SK (neg) | -0.004 | 0.042 | 2904.3 | -0.11 | 0.915 |
|  | Cue SK (pos) | -0.046 | 0.042 | 2905.3 | -1.11 | 0.269 |
|  | Cue mean x var | 0.047 | 0.029 | 2912.8 | 1.58 | 0.114 |
|  | Stim int x cue mean | -0.02 | 0.029 | 2903.8 | -0.69 | 0.49 |
| Right NAc shell | Stimulus Intensity | 0.026 | 0.03 | 2907.9 | 0.87 | 0.384 |
|  | Cue mean | 0.043 | 0.031 | 304.7 | 1.39 | 0.166 |
|  | Cue var | -0.052 | 0.03 | 2908.8 | -1.73 | 0.084 |
|  | Cue SK (neg) | 0.038 | 0.043 | 2907.7 | 0.88 | 0.378 |
|  | Cue SK (pos) | -0.005 | 0.043 | 2908.9 | -0.11 | 0.909 |
|  | Cue mean x var | 0.039 | 0.03 | 2915.3 | 1.31 | 0.192 |
|  | Stim int x cue mean | 0.029 | 0.03 | 2907.1 | 0.95 | 0.344 |
| Left NAc core | Stimulus Intensity | 0.033 | 0.03 | 2947.9 | 1.1 | 0.273 |
|  | Cue mean | 0.054 | 0.03 | 2948.4 | 1.81 | 0.07 |
|  | **Cue var** | **-0.104** | **0.03** | **2949.7** | **-3.47** | **0.001** |
|  | Cue SK (neg) | 0.019 | 0.042 | 2948.6 | 0.44 | 0.66 |
|  | Cue SK (pos) | -0.031 | 0.042 | 2949.1 | -0.73 | 0.465 |
|  | **Cue mean x var** | **0.073** | **0.03** | **2948.3** | **2.43** | **0.015** |
|  | Stim int x cue mean | -0.006 | 0.03 | 2948.6 | -0.2 | 0.841 |
| Right NAc core | Stimulus Intensity | 0.045 | 0.03 | 2948.3 | 1.51 | 0.131 |
|  | Cue mean | 0.021 | 0.03 | 2948.8 | 0.69 | 0.49 |
|  | Cue var | -0.02 | 0.03 | 2950.1 | -0.67 | 0.5 |
|  | Cue SK (neg) | 0.032 | 0.042 | 2948.9 | 0.76 | 0.445 |
|  | Cue SK (pos) | 0.004 | 0.042 | 2949.5 | 0.09 | 0.929 |
|  | Cue mean x var | 0.045 | 0.03 | 2948.6 | 1.5 | 0.133 |
|  | Stim int x cue mean | -0.002 | 0.03 | 2949 | -0.06 | 0.956 |

**Supplementary Table 8.** Statistics for the effects on ROIs related to higher level visual processing. Abbreviations: var = variance; SK = skewness; neg = negative (vs. symmetric); pos = positive (vs. symmetric); stim int = stimulus intensity; IPS = intraparietal sulcus.

| Region | Effect | Estimate | SE | DF | t | p |
| --- | --- | --- | --- | --- | --- | --- |
| Left IPS | Stimulus Intensity | 0.012 | 0.029 | 2958.7 | 0.40 | 0.691 |
|  | Cue mean | -0.054 | 0.029 | 2958.2 | -1.84 | 0.066 |
|  | Cue var | -0.021 | 0.029 | 2958.8 | -0.71 | 0.479 |
|  | Cue SK (neg) | -0.030 | 0.041 | 2958.6 | -0.73 | 0.468 |
|  | Cue SK (pos) | -0.004 | 0.041 | 2958.5 | -0.11 | 0.915 |
|  | Cue mean x var | 0.029 | 0.029 | 2958.1 | 0.99 | 0.325 |
|  | Stim int x cue mean | -0.013 | 0.029 | 2958.1 | -0.43 | 0.664 |
| Right IPS | Stimulus Intensity | 0.004 | 0.029 | 2956.7 | 0.14 | 0.891 |
|  | Cue mean | -0.052 | 0.029 | 2956.2 | -1.79 | 0.074 |
|  | Cue var | -0.036 | 0.029 | 2956.8 | -1.24 | 0.216 |
|  | Cue SK (neg) | -0.013 | 0.042 | 2956.6 | -0.31 | 0.753 |
|  | Cue SK (pos) | 0.000 | 0.041 | 2956.5 | -0.01 | 0.991 |
|  | Cue mean x var | 0.054 | 0.029 | 2956.2 | 1.86 | 0.064 |
|  | Stim int x cue mean | -0.015 | 0.029 | 2956.3 | -0.52 | 0.605 |

**Supplementary Table 9.** Statistics for the effects on other pain processing ROIs. Abbreviations: var = variance; SK = skewness; neg = negative (vs. symmetric); pos = positive (vs. symmetric); stim int = stimulus intensity; aIns = anterior insula. Significant effects (uncorrected *p* < .05) are marked in bold.

| Region | Effect | Estimate | SE | DF | t | p |
| --- | --- | --- | --- | --- | --- | --- |
| Right aIns | **Stimulus Intensity** | **0.076** | **0.028** | **2904.4** | **2.68** | **0.007** |
|  | Cue mean | 0.018 | 0.029 | 295.0 | 0.61 | 0.540 |
|  | Cue var | 0.008 | 0.028 | 2904.1 | 0.29 | 0.773 |
|  | Cue SK (neg) | 0.007 | 0.040 | 2903.7 | 0.18 | 0.858 |
|  | Cue SK (pos) | 0.071 | 0.040 | 2904.8 | 1.78 | 0.075 |
|  | Cue mean x var | 0.002 | 0.028 | 2912.1 | 0.07 | 0.942 |
|  | Stim int x cue mean | -0.040 | 0.028 | 2903.2 | -1.41 | 0.158 |
| Left aIns | **Stimulus Intensity** | **0.093** | **0.029** | **2948.3** | **3.21** | **0.001** |
|  | Cue mean | -0.016 | 0.029 | 2948.7 | -0.54 | 0.588 |
|  | Cue var | 0.002 | 0.029 | 2949.6 | 0.06 | 0.951 |
|  | Cue SK (neg) | 0.024 | 0.041 | 2948.8 | 0.57 | 0.567 |
|  | Cue SK (pos) | 0.045 | 0.041 | 2949.1 | 1.08 | 0.278 |
|  | Cue mean x var | -0.007 | 0.029 | 2948.6 | -0.23 | 0.820 |
|  | Stim int x cue mean | -0.010 | 0.029 | 2948.9 | -0.33 | 0.739 |
| Right S1 hand | **Stimulus Intensity** | **0.117** | **0.026** | **2948.2** | **4.49** | **0.000** |
|  | Cue mean | -0.007 | 0.026 | 2948.4 | -0.27 | 0.788 |
|  | Cue var | -0.009 | 0.026 | 2948.8 | -0.33 | 0.738 |
|  | Cue SK (neg) | 0.015 | 0.037 | 2948.4 | 0.40 | 0.689 |
|  | Cue SK (pos) | 0.036 | 0.037 | 2948.6 | 0.99 | 0.324 |
|  | Cue mean x var | 0.018 | 0.026 | 2948.4 | 0.67 | 0.500 |
|  | Stim int x cue mean | -0.010 | 0.026 | 2948.5 | -0.37 | 0.709 |
| Left S1 hand | **Stimulus Intensity** | **0.133** | **0.027** | **2908.5** | **5.00** | **0.000** |
|  | Cue mean | -0.033 | 0.027 | 357.6 | -1.21 | 0.225 |
|  | Cue var | -0.012 | 0.027 | 2907.8 | -0.46 | 0.645 |
|  | Cue SK (neg) | 0.034 | 0.038 | 2907.6 | 0.90 | 0.370 |
|  | Cue SK (pos) | 0.024 | 0.038 | 2908.5 | 0.63 | 0.527 |
|  | Cue mean x var | -0.020 | 0.027 | 2916.3 | -0.76 | 0.445 |
|  | Stim int x cue mean | -0.009 | 0.027 | 2907.2 | -0.34 | 0.736 |

**Supplementary Table 10.** Statistics for the effects on other visual processing ROIs. Abbreviations: var = variance; SK = skewness; neg = negative (vs. symmetric); pos = positive (vs. symmetric); stim int = stimulus intensity. Significant effects (uncorrected *p* < .05) are marked in bold.

| Region | Effect | Estimate | SE | DF | t | p |
| --- | --- | --- | --- | --- | --- | --- |
| Right V3A | Stimulus Intensity | 0.041 | 0.027 | 2958 | 1.54 | 0.124 |
|  | Cue mean | -0.014 | 0.027 | 2957.7 | -0.54 | 0.590 |
|  | Cue var | 0.014 | 0.027 | 2958 | 0.54 | 0.590 |
|  | **Cue SK (neg)** | **-0.087** | **0.038** | **2957.9** | **-2.30** | **0.021** |
|  | Cue SK (pos) | -0.040 | 0.038 | 2957.9 | -1.05 | 0.292 |
|  | Cue mean x var | 0.000 | 0.027 | 2957.7 | 0.02 | 0.987 |
|  | Stim int x cue mean | -0.003 | 0.027 | 2957.7 | -0.11 | 0.911 |
| Left V3A | Stimulus Intensity | 0.043 | 0.028 | 2955.5 | 1.57 | 0.115 |
|  | Cue mean | -0.012 | 0.028 | 2955.2 | -0.45 | 0.653 |
|  | Cue var | -0.003 | 0.028 | 2955.5 | -0.10 | 0.920 |
|  | **Cue SK (neg)** | **-0.101** | **0.039** | **2955.4** | **-2.57** | **0.010** |
|  | Cue SK (pos) | -0.052 | 0.039 | 2955.3 | -1.33 | 0.184 |
|  | Cue mean x var | -0.022 | 0.028 | 2955.1 | -0.82 | 0.415 |
|  | Stim int x cue mean | -0.004 | 0.028 | 2955.2 | -0.13 | 0.898 |
| Right V3B | Stimulus Intensity | 0.033 | 0.029 | 2961.9 | 1.13 | 0.260 |
|  | Cue mean | -0.046 | 0.029 | 2961.4 | -1.58 | 0.114 |
|  | Cue var | -0.013 | 0.029 | 2962 | -0.44 | 0.657 |
|  | Cue SK (neg) | -0.046 | 0.041 | 2961.8 | -1.13 | 0.260 |
|  | Cue SK (pos) | -0.029 | 0.041 | 2961.7 | -0.71 | 0.476 |
|  | Cue mean x var | 0.040 | 0.029 | 2961.4 | 1.39 | 0.164 |
|  | Stim int x cue mean | 0.012 | 0.029 | 2961.4 | 0.42 | 0.672 |
| Left V3B | Stimulus Intensity | 0.007 | 0.028 | 2960.9 | 0.24 | 0.811 |
|  | Cue mean | -0.019 | 0.028 | 2960.5 | -0.68 | 0.494 |
|  | Cue var | -0.024 | 0.028 | 2960.8 | -0.88 | 0.379 |
|  | Cue SK (neg) | -0.055 | 0.039 | 2960.7 | -1.40 | 0.163 |
|  | Cue SK (pos) | -0.023 | 0.039 | 2960.6 | -0.58 | 0.562 |
|  | Cue mean x var | -0.003 | 0.028 | 2960.4 | -0.11 | 0.911 |
|  | Stim int x cue mean | 0.018 | 0.028 | 2960.5 | 0.64 | 0.523 |
| Right V3CD | Stimulus Intensity | 0.032 | 0.026 | 2946.2 | 1.23 | 0.219 |
|  | Cue mean | -0.022 | 0.026 | 2946 | -0.85 | 0.393 |
|  | Cue var | -0.046 | 0.026 | 2946.2 | -1.76 | 0.079 |
|  | Cue SK (neg) | -0.064 | 0.037 | 2946.2 | -1.74 | 0.083 |
|  | Cue SK (pos) | -0.052 | 0.037 | 2946.1 | -1.41 | 0.160 |
|  | Cue mean x var | 0.031 | 0.026 | 2946 | 1.19 | 0.234 |
|  | Stim int x cue mean | 0.021 | 0.026 | 2946 | 0.81 | 0.416 |
| Left V3CD | Stimulus Intensity | 0.001 | 0.026 | 2960.6 | 0.06 | 0.955 |
|  | Cue mean | -0.015 | 0.026 | 2960.3 | -0.56 | 0.575 |
|  | Cue var | -0.038 | 0.026 | 2960.6 | -1.48 | 0.139 |
|  | Cue SK (neg) | -0.020 | 0.037 | 2960.5 | -0.54 | 0.588 |
|  | Cue SK (pos) | -0.035 | 0.037 | 2960.4 | -0.94 | 0.346 |
|  | Cue mean x var | 0.013 | 0.026 | 2960.3 | 0.51 | 0.609 |
|  | Stim int x cue mean | -0.008 | 0.026 | 2960.3 | -0.30 | 0.765 |
| Right V4t | Stimulus Intensity | 0.001 | 0.029 | 2952.9 | 0.04 | 0.968 |
|  | Cue mean | 0.008 | 0.029 | 2952.4 | 0.26 | 0.792 |
|  | Cue var | -0.021 | 0.029 | 2952.9 | -0.73 | 0.466 |
|  | Cue SK (neg) | -0.062 | 0.040 | 2952.7 | -1.52 | 0.128 |
|  | Cue SK (pos) | -0.042 | 0.040 | 2952.6 | -1.05 | 0.293 |
|  | Cue mean x var | 0.048 | 0.029 | 2952.4 | 1.67 | 0.095 |
|  | Stim int x cue mean | 0.002 | 0.029 | 2952.4 | 0.05 | 0.957 |
| Left V4t | Stimulus Intensity | -0.002 | 0.027 | 2961.3 | -0.07 | 0.945 |
|  | Cue mean | -0.022 | 0.027 | 2961.1 | -0.84 | 0.398 |
|  | Cue var | -0.041 | 0.027 | 2961.4 | -1.54 | 0.125 |
|  | Cue SK (neg) | -0.062 | 0.038 | 2961.3 | -1.64 | 0.101 |
|  | Cue SK (pos) | -0.056 | 0.038 | 2961.2 | -1.48 | 0.138 |
|  | Cue mean x var | 0.020 | 0.027 | 2961.1 | 0.73 | 0.463 |
|  | Stim int x cue mean | 0.026 | 0.027 | 2961.1 | 0.97 | 0.334 |
| Right<br>ventro-medial<br>V1 | Stimulus Intensity | 0.023 | 0.030 | 2955.6 | 0.79 | 0.432 |
|  | Cue mean | 0.020 | 0.030 | 2954.8 | 0.66 | 0.510 |
|  | Cue var | -0.013 | 0.030 | 2955.6 | -0.43 | 0.668 |
|  | Cue SK (neg) | -0.035 | 0.042 | 2955.4 | -0.82 | 0.413 |
|  | Cue SK (pos) | -0.012 | 0.042 | 2955.2 | -0.28 | 0.780 |
|  | Cue mean x var | -0.018 | 0.030 | 2954.7 | -0.61 | 0.542 |
|  | Stim int x cue mean | 0.005 | 0.030 | 2954.9 | 0.18 | 0.856 |
| Left<br>ventro-medial<br>V1 | Stimulus Intensity | 0.007 | 0.030 | 2915.6 | 0.25 | 0.806 |
|  | Cue mean | -0.017 | 0.031 | 330.5 | -0.55 | 0.585 |
|  | Cue var | 0.033 | 0.030 | 2915 | 1.09 | 0.275 |
|  | Cue SK (neg) | -0.026 | 0.043 | 2915.2 | -0.62 | 0.537 |
|  | Cue SK (pos) | 0.005 | 0.043 | 2914.6 | 0.12 | 0.903 |
|  | Cue mean x var | -0.017 | 0.030 | 2916.3 | -0.56 | 0.577 |
|  | Stim int x cue mean | 0.014 | 0.030 | 2917.3 | 0.48 | 0.632 |
| Right<br>ventro-medial<br>V2 | Stimulus Intensity | 0.018 | 0.030 | 234.8 | 0.58 | 0.564 |
|  | Cue mean | -0.005 | 0.030 | 266.1 | -0.18 | 0.857 |
|  | Cue var | 0.013 | 0.029 | 2875.1 | 0.45 | 0.650 |
|  | Cue SK (neg) | -0.068 | 0.040 | 2874.7 | -1.68 | 0.094 |
|  | Cue SK (pos) | -0.039 | 0.040 | 2880.5 | -0.96 | 0.339 |
|  | Cue mean x var | -0.026 | 0.029 | 2875.7 | -0.91 | 0.362 |
|  | Stim int x cue mean | -0.005 | 0.029 | 2877.1 | -0.16 | 0.874 |
| Left<br>ventro-medial<br>V2 | Stimulus Intensity | -0.021 | 0.029 | 2959.2 | -0.73 | 0.466 |
|  | Cue mean | -0.026 | 0.029 | 2958.7 | -0.91 | 0.360 |
|  | Cue var | -0.020 | 0.029 | 2959.2 | -0.70 | 0.486 |
|  | Cue SK (neg) | -0.006 | 0.041 | 2959 | -0.16 | 0.876 |
|  | Cue SK (pos) | -0.024 | 0.041 | 2958.9 | -0.58 | 0.564 |
|  | Cue mean x var | -0.011 | 0.029 | 2958.6 | -0.38 | 0.702 |
|  | Stim int x cue mean | -0.008 | 0.029 | 2958.7 | -0.26 | 0.793 |
| Right<br>ventro-medial<br>V3 | Stimulus Intensity | 0.033 | 0.028 | 2911 | 1.16 | 0.247 |
|  | Cue mean | -0.014 | 0.029 | 319.8 | -0.48 | 0.629 |
|  | Cue var | 0.016 | 0.028 | 2910.1 | 0.55 | 0.582 |
|  | Cue SK (neg) | -0.078 | 0.040 | 2910.3 | -1.93 | 0.053 |
|  | Cue SK (pos) | -0.046 | 0.040 | 2910.5 | -1.15 | 0.249 |
|  | Cue mean x var | 0.000 | 0.028 | 2912.5 | 0.02 | 0.987 |
|  | Stim int x cue mean | -0.014 | 0.028 | 2913 | -0.49 | 0.624 |
| Left<br>ventro-medial<br>V3 | Stimulus Intensity | 0.027 | 0.028 | 2962.7 | 0.98 | 0.326 |
|  | Cue mean | -0.027 | 0.028 | 2962.5 | -0.97 | 0.330 |
|  | Cue var | -0.012 | 0.028 | 2962.8 | -0.45 | 0.651 |
|  | Cue SK (neg) | -0.057 | 0.039 | 2962.8 | -1.45 | 0.147 |
|  | Cue SK (pos) | -0.033 | 0.039 | 2962.6 | -0.84 | 0.403 |
|  | Cue mean x var | 0.006 | 0.028 | 2962.4 | 0.23 | 0.819 |
|  | Stim int x cue mean | -0.019 | 0.028 | 2962.5 | -0.70 | 0.482 |

